# UPP2: Fast and Accurate Alignment Estimation of Datasets with Fragmentary Sequences

**DOI:** 10.1101/2022.02.26.482099

**Authors:** Minhyuk Park, Stefan Ivanovic, Gillian Chu, Chengze Shen, Tandy Warnow

## Abstract

**Motivation:** Multiple sequence alignment (MSA) is a basic step in many bioinformatics pipelines. However, achieving highly accurate alignments on large datasets, especially those with sequence length heterogeneity, is a challenging task. UPP (Ultra-large multiple sequence alignment using Phylogeny-aware Profiles) is a method for MSA estimation that builds an ensemble of Hidden Markov Models (eHMM) to represent an estimated alignment on the full length sequences in the input, and then adds the remaining sequences into the alignment using selected HMMs in the ensemble. Although UPP provides good accuracy, it is computationally intensive on large datasets.

**Results:** We present UPP2, a direct improvement on UPP. The main advance is a fast technique for selecting HMMs in the ensemble that allows us to achieve the same accuracy as UPP but with greatly reduced runtime. We show UPP2 produces more accurate alignments compared to leading MSA methods on datasets exhibiting substantial sequence length heterogeneity, and is among the most accurate otherwise.

**Availability:** https://github.com/gillichu/sepp

## 1 Introduction

Multiple sequence alignment is a fundamental bioinformatics task, and producing accurate alignments can have profound impact in many downstream analyses such as phylogeny inference (Morrison and Ellis, 1997), detection of adaptive evolution (Blackburne and Whelan, 2013), or protein structure and function inference (Bork and Koonin, 1998; Ju *et al*., 2021).

Because of the significant interest in alignment estimation, many alignment methods have been developed (e.g., MUSCLE (Edgar, 2004), PRANK (Löytynoja and Goldman, 2005), BAli-Phy (Suchard and Redelings, 2006), Clustal Omega (Sievers *et al*., 2011), MAFFT (Katoh and Standley, 2013), PASTA (Mirarab *et al*. 2015), MAGUS (Smirnov and Warnow, 2021a), and regressive T-COFFEE Garriga *et al*., 2019)). However, accurate alignment is still challenging under some conditions. For example, large datasets (with many thousands of sequences) can be difficult to align with high accuracy and also present substantial computational challenges (Nguyen *et al*., 2015; Mirarab *et al*., 2015, Smirnov, 2021). The difficulty in aligning datasets that are highly heterogeneous due to high rates of evolution has also been documented (Liu *et al*., 2009), but several methods (largely employing divide-and-conquer) have been able to achieve good accuracy in such conditions (e.g., PASTA (Mirarab *et al*., 2015) and MAGUS (Smirnov and Warnow, 2021a)). Sequence length heterogeneity (Figure 1) introduces another challenge for alignment estimation, and one that is relatively less studied (Nguyen *et al*., 2015. Shen *et al*., 2021).

**Figure 1:**
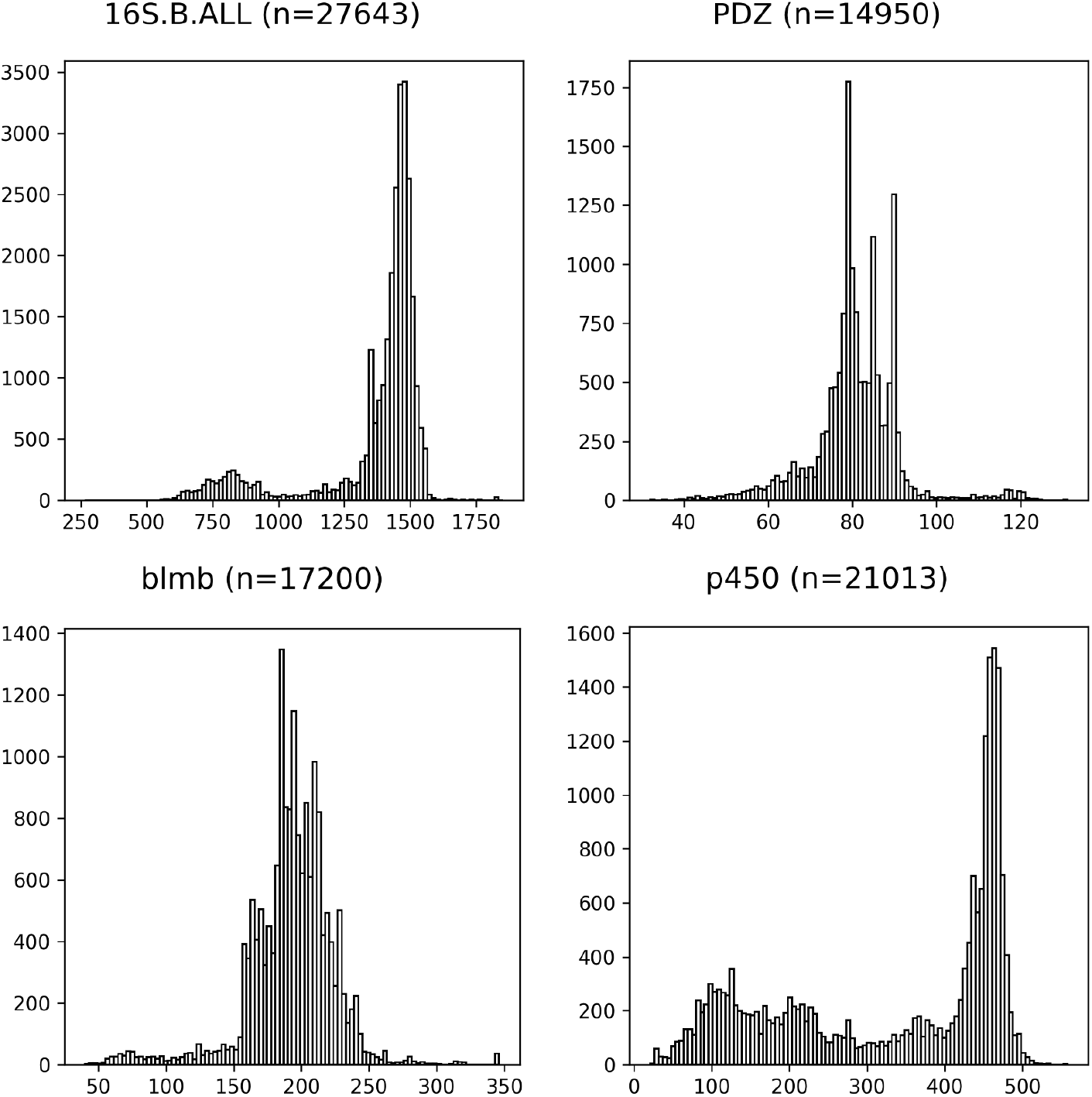
Histograms of sequence lengths in biological datasets. CRW 16S.B.ALL is from Cannone *et al*. (2002) and the datasets in the other three panels are from the Homfam collection Blackshields *et al*. (2010), which consists of HOMSTRAD reference sequences with Pfam sequences from the same domain.

UPP (Ultra-large alignments using Phylogeny-aware Profiles) (Nguyen *et al*., 2015)) is a multiple sequence alignment method that was specifically designed to provide good accuracy on datasets with substantial sequence length heterogeneity, while maintaining scalability on large datasets. UPP operates in three basic stages: first, it extracts and aligns a subset of the sequences it deems to be full-length; second, it builds an ensemble of Hidden Markov Models (HMMs) (Durbin *et al*., 1998) on the alignment of the selected full-length sequences; and third, it uses the ensemble to align the remaining sequences. Thus, UPP uses a combination of global MSA methods (to align the backbone sequences, which are full-length) and local MSA methods (to add the remaining sequences into the backbone alignment, which include sequences that are short).

This third step is often the bottleneck in terms of runtime. Specifically, for each additional sequence that needs to be aligned, the HMM with the highest bit-score is selected from the ensemble and is used to add the sequence into the alignment. By design, the first two steps are reasonably fast, but the third step requires an all-against-all comparison of the remaining sequences against the HMMs in the ensemble. Thus, the runtime of UPP can be prohibitively high when there are many sequences that are not full-length and when the ensemble contains many HMMs.

In the last year, modifications to UPP to improve its accuracy and theoretical foundation have been explored. The default for UPP as provided in the github site uses PASTA to align the backbone sequences. However, Shen *et al*. (2021) showed that alignment accuracy was improved by using MAGUS instead of PASTA to compute the backbone alignment. Another potential weakness in the original UPP approach is the use of the bit score to select the single HMM to align the query sequence. A bit-score represents the log likelihood ratio of a query sequence being emitted by an HMM to the likelihood of a query sequence being emitted by a null HMM. However, the bit-score does not correspond to the probability that the query sequence is generated by the selected HMM from the ensemble, as this specific question depends also on the number of sequences used to build the HMM as well as the ensemble of HMMs that has been constructed. To address this, a modification to the use of bit-scores, called “adjusted bitscores”, was presented in Shen *et al*. (2022). Under the assumption that exactly one of the HMMs in the ensemble generated the query sequence, adjusted bitscores can be interpreted as probabilities that the given HMM generates the query sequence (Shen *et al*., 2022). The Supplementary Materials (Section S2) provides the formula for the adjusted bit-score, its derivation, and additional discussion.

Although the version in the UPP Github site still uses PASTA for the backbone and selects the best HMM based on raw bit-scores, based on these two studies, the current recommended setting for UPP uses MAGUS for the backbone alignment and selects the “best” HMM from the ensemble based on the adjusted bit-score.

These modifications aimed to improve accuracy rather than runtime, and UPP has remained computationally intensive as a result of its all-against-all algorithmic design. Here we present UPP2, a modification to UPP that is designed to reduce its runtime and improve its scalability to large sequence datasets. The main modification we use is a replacement of the all-against-all comparison of query sequences and HMMs by a much smaller number of comparisons, so that each query sequence is scored against a logarithmic number of HMMs instead of against all the HMMs. As we will show, this change reduces the runtime, sometimes dramatically, without hurting accuracy.

## 2 UPP2

### 2.1 The UPP three-stage pipeline

In the first stage, it computes a backbone alignment and backbone tree on a subset of the input sequences, in the second stage it builds an ensemble of profile HMMs on the backbone alignment, and in the third stage it uses the ensemble to add all the remaining sequences into the backbone alignment using commands from HMMER (Eddy, 2011) (hmmbuild, hmmsearch, and hmmalign). Here we provide some additional details.

For Stage 1, by default, UPP will select up to 1000 sequences to include in its backbone, and these sequences are selected at random from the set of sequences within 25% in length of the median length sequence. The alignment is built using a selected “base method”, with PASTA the original technique and now MAGUS the recommended technique. Our own studies have suggested that larger backbones may improve final alignment accuracy; hence, using 10,000 sequences for the backbone on large datasets (e.g., with at least 25,000 sequences) is the approach that we follow in this study.

For Stage 2, UPP computes a set of subset alignments by hierarchically decomposing the backbone tree at a centroid edge (i.e., an edge that splits the leaf set into two sets of roughly equal sizes) until all the subtrees are at most size *z*, where *z* is an input to UPP. UPP builds an HMM on each set created during this decomposition, including the full set, thus producing a collection of HMMs that we refer to as the “ensemble of HMMs” (eHMM) for the backbone alignment. In the initial version of UPP, *z* was set to 10. Some studies (Mirarab *et al*. that developed eHMMs for other purposes have suggested that smaller values (e.g., *z* = 2) might improve accuracy, but a more recent study exploring this question for alignment estimation Shen *et al*. (2021) has found otherwise.

For Stage 3, UPP adds every additional sequence (i.e., ones that are not in the backbone) into the backbone alignment. These additional sequences are referred to as “query sequences” and are added as follows. For each query sequence, *hmmsearch* is used to find the HMM that returns the highest bit-score (the original setting) or the highest adjusted bit-score (the current recommendation). Then, each query sequence is added into the subset alignment used to construct the selected HMM using *hmmalign*. Since the subset alignments are induced by the backbone alignment, this also means the query sequence can be added into the backbone alignment as well. The addition of the query sequence into the backbone alignment defines an “extended alignment”. The extended alignments from the different query sequences are merged together using transitivity, thus producing a final alignment containing all the sequences.

### 2.2 UPP2: Modifying Stage 3 to improve speed

In Stage 3, each query sequence picks a best HMM (based on the bit-score or the adjusted bit-score) and then that HMM is used to add the query sequence into the backbone alignment. Due to its hierarchical decomposition strategy, UPP produces many HMMs, all of which have to be compared against every single query sequence. This quickly presents scalability issues in several cases: as the size of backbone increases, as *z* (which defines the decomposition stopping rule) decreases, or as the number of query sequences increases. We propose two strategies (“Hierarchical” and “EarlyStop”) based on this hierarchical decomposition strategy to speed up the search: Hierarchical and EarlyStop (Figure 2). We denote UPP with these strategies using the notation “UPP+Hierarchical” or “UPP+EarlyStop”.

**Figure 2:**
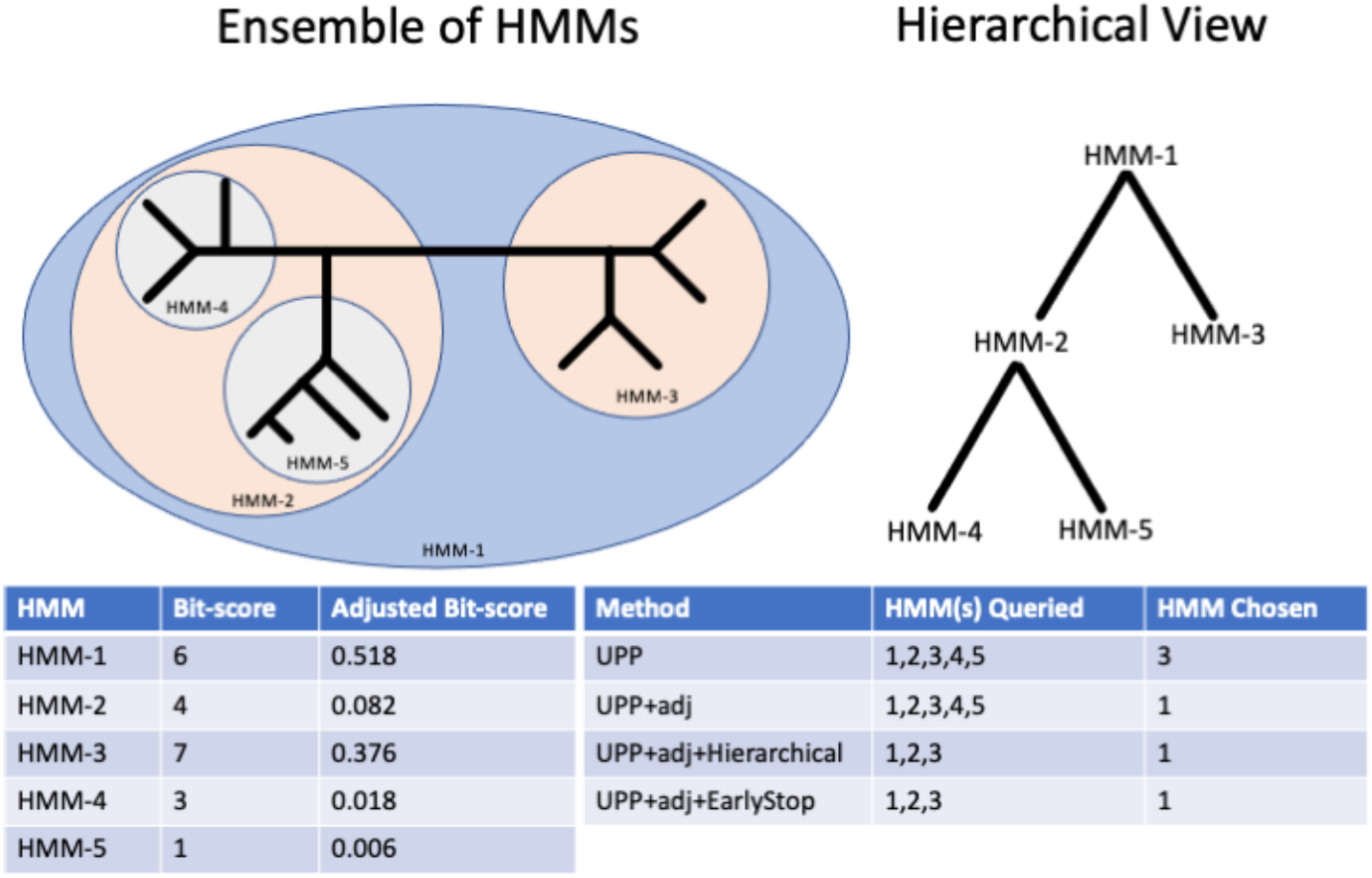
UPP and UPP2 Search Strategies (Toy Example) Here we show a sample ensemble of HMMs and how different search strategies pick different HMMs within the ensemble. UPP and UPP+adj by default search through every HMM but use different criteria (raw bit-scores or adjusted bit-scores, respectively). UPP will choose HMM-3 since HMM-3 has the highest bit-score while UPP+adj will choose HMM-1 since HMM-1 has the highest adjusted bit-score. UPP+adj+Hierarchical will start at HMM-1 and descend down the subtree with the highest adjusted bit-score. UPP+adj+EarlyStop will descend down the subtree with the highest adjusted bit-score and stop once all immediate children HMMs have worse adjusted bit-scores than the current best HMM.

#### Hierarchical

Stage 2 defines a hierarchy of HMMs based on their sequence sets, so that the set of HMMs forms a rooted tree. Here we describe how the Hierarchical Search strategy operates, using adjusted bit-scores. To select an HMM for a given query sequence *q*, we start at the root HMM and we compute its adjusted bit-score given *q*. We then evaluate the children HMMs and descend down into the subtree that has the larger adjusted bit-score (randomly selecting one in the case of a tie). The process continues until a leaf HMM is reached. The HMM with the largest adjusted bit-score (i.e., the HMM deemed the most likely to have emitted the query sequence) encountered during the traversal then becomes selected HMM for the query sequence. Note that this strategy evaluates at most two HMMs per level in the tree. In the case of a tie, the HMM that comes first in a pre-order traversal is chosen.

#### EarlyStop

We follow the same basic strategy as Hierarchical. However, the process stops descending down the subtree in the hierarchical search process if both of the two children HMMs have lower adjusted bit-scores, and are therefore considered less likely to have emitted the query sequence, than the current best HMM (hence the name “EarlyStop”).

## 3 Experimental Study

### Overview

We performed two experiments, one for designing UPP2 (Experiment 1) and one for evaluating UPP2 in comparison to leading alignment methods (Experiment 2). Experiment 1 was performed on a small set of “training datasets” and Experiment 2 was performed on a larger set of “testing datasets”. Methods were evaluated for alignment error and runtime.

#### Alignment methods

We evaluated variants of UPP2 that differ in terms of the backbone alignment method (PASTA or MAGUS), the use of raw or adjusted bit-scores, the stopping condition (i.e., how *z* is set), and whether the all-against-all comparisons are performed or one of the two faster search strategies is used. Recall that the original version of UPP uses PASTA backbones, raw bit-scores, sets *z* = 10, and performs all-against-all comparisons to find the best HMM for each query sequence. We explore the following variants of UPP2 (indicating how they differ from the original version of UPP below). Each of these versions can have either PASTA or MAGUS backbones, as indicated in parentheses.

- *UPP+adj*: UPP+adj differs from the original UPP by using adjusted bit-scores.
- *UPP+adj+Hierarchical*: Identical to UPP+adj except that it uses the Hierarchical search strategy.
- *UPP+adj+EarlyStop*: Identical to UPP+adj except that it uses the EarlyStop search strategy.

We also evaluated the following leading alignment methods:

- **MUSCLE** (3.8.31), limited to 2 iterations.
- **Clustal Omega** (1.2.4), used in its default mode.
- **T-COFFEE** (13.45.0.4846264), used in the default regressive mode.
- **MAGUS**, (Commit on 4/5/21, commit ID in supplement) used in its default mode, which is with recursion in the newest version of MAGUS (Smirnov and Warnow, 2021a; Smirnov, 2021).
- **PASTA** (v1.9.0), used in its default mode.
- **MAFFT** (7.487), with the *linsi* mode used for datasets of size at most 1000 and the *auto* mode used for larger datasets.
- **UPP** (through appropriate settings of the algorithmic parameters within the UPP2 code).

#### Datasets

We used both biological and simulated datasets (both nucleotides and proteins) for the experiments, separating them into the training datasets (used in Experiment 1) and the testing datasets (used in Experiment 2). We had fragmentary versions of the datasets; for these datasets, the suffix “HF” denotes high fragmentary datasets, which are constructed by taking the original dataset and making half of the sequences 25% of the original median sequence length. The fragmentation process is explained in full detail in Smirnov and Warnow (2021b). The empirical statistics (i.e., number of sequences, average sequence length, percent of the reference alignment occupied by gaps, and average and maximum p-distance) for these datasets are provided in the Supplementary Materials Table S1 (for the nucleotide datasets) and Table S2 (for the protein datasets) The sequence length histograms for the biological datasets are provided in Supplementary Figures S1 to S13.

The ROSE simulated datasets, introduced in Liu *et al*. (2009), are 1000-sequence datasets with varying gap lengths, which are denoted by “S” for short gap lengths, “M” for medium gap lengths, and “L” for long gap lengths. We used 1000S1 through 1000S5, 1000M1 through 1000M5, and 1000L1 through 1000L5 as well as their high fragmentary counterparts 1000S1-HF through 1000S5-HF, 1000M1-HF through 1000M5-HF, and 1000L1-HF through 1000L5-HF. 1000M1 and 1000M1-HF were using for training while the other model conditions were reserved for testing. These datasets range in alignment difficulty as a result of the evolutionary rate, as reflected in their average p-distance (i.e., average normalized Hamming distance between any two sequences). The most difficult model conditions are 1000M1, 1000S1, and 1000L1, and the easiest ones are 1000S5, 1000M4, 1000M5, and 1000L5. Comparing the p-distances, we see that the difficult model conditions have average p-distances at least 69% and the easiest model conditions have average p-distances below 50%.

RNASim datasets are created by sampling from the RNASim million-sequence dataset, originally created by Guo *et al*. (2009) and studied in Mirarab *et al*. (2015). We used the same 1000-sequence RNASim datasets as Smirnov and Warnow (2021b), which are published at https://doi.org/10.5061/dryad.95x69p8h8; these are the RNASim1000 and RNASim1000-HF. RNASim1000 and RNASim1000-HF datasets. The RNASim datasets have average p-distance of 41%.

The 16S datasets are biological datasets based on the 16S gene. The 16S.B.ALL, 16S.3, and 16S.T datasets were used in the study by Liu *et al*. (2011). These datasets have reference alignments based on secondary structure (Cannone *et al*., 2002) and are available at https://sites.google.com/eng.ucsd.edu/datasets/alignment/16s23s.

The Homfam datasets are biological datasets created by combining small numbers of HOMSTRAD reference sequences with Pfam sequences from the same domain, originally compiled by Blackshields *et* al. (2010). These were used in the study by Mirarab *et al*. (2015) and are available at https://sites.google.com/eng.ucsd.edu/datasets/alignment/pastaupp. We picked the 10 largest Homfam datasets for our study. These datasets only have reference alignments on the HOMSTRAD sequences, so that alignment error can only be evaluated on a small subset of sequences for each dataset.

For the training datasets (Experiment 1), we used two model conditions, 1000M1 (one of the model conditions from the ROSE simulated datasets with high rates of evolution) and RNASim1000. For each model condition, we explored full-length versions and HF versions. We used the remaining datasets as the testing datasets (Experiment 2).

#### Computational Resources

Experiment 1 was done on Blue Waters (Bode *et al*., 2013). In Experiment 2a, UPP(MAGUS)+adj and UPP2 were run on the Illinois Campus Cluster while all other methods were run on Blue Waters. In Experiment 2b, UPP(MAGUS)+adj, UPP2, and MAFFT were run on the Illinois Campus Cluster while all other methods were run on Blue Waters. In Experiment 2c, all analyses were run on the Illinois Campus Cluster. We only show runtime comparisons when the methods were run on the same system under the same conditions. All methods were limited to a maximum of 7 days of wall-time and a maximum of 256 GB of RAM. MUSCLE does not have a multi-threaded version and was unable to take advantage of the core count. Some failures to complete due to limitations of time and/or memory occurred; these are reported in detail in the Supplementary Materials Section S6.

#### Alignment Error

We used FastSP (1.7.1) (Mirarab and Warnow, 2011) for calculating SPFN and SPFP rates of estimated alignments relative to the reference alignments, defined as follows. SPFN refers to “sum-of-pairs false negatives”, and is the number of the pairwise homologies found in the reference alignment but not in the estimated alignment, while SPFP refers to “sum-of-pairs false positives” and is the number of pairwise homologies found in the estimated alignment but not in the reference alignment. These are normalized by the number of homologies in the reference alignment or estimated alignment, respectively, to produce the SPFN and SPFP error rates.

#### Experiment 1 Overview

Experiment 1 explored the impact of decomposition size (i.e., value for *z*), use of adjusted bit-scores, and new search strategies (Hierarchical and EarlyStop) for UPP2. We additionally evaluate the impact of using MAGUS backbone alignments instead of PASTA backbone alignments for use in UPP2. We used 1000M1, RNASim1000, 1000M1-HF, and RNASim1000-HF. For 1000M1 and RNASim1000, 500 full length backbone sequences were selected by UPP. 1000M1-HF and RNASim1000-HF, the full-length sequences are easily identified, and that information is provided to UPP.

#### Experiment 2 Overview

We used Experiment 1 to specify to set the algorithmic parameters for the approach and refer to this variant as “UPP2”. Experiment 2 compared UPP2, UPP(MAGUS)+adj, MAGUS, PASTA, MAFFT (using *linsi* for datasets with at most 1000 sequences and *auto* otherwise), Clustal Omega, regressive T-COFFEE, and MUSCLE. Experiment 2a examined results on simulated datasets with fragmentary sequences, Experiment 2b examined results on 16S biological datasets, and Experiment 2c examined results on the Homfam biological datasets. We used the testing datasets for this experiment. The default number of sequences in the backbone alignments depends on the dataset size: for these datasets with more than 25,000 sequences we used 10,000 sequences for the backbone, and the default for the others was 1000 sequences. In Experiment 2c (on the Homfam datasets), we also explored the impact of even larger backbones, where we used all “full-length” sequences. The definition of full-length for the Homfam datasets was 25% within the length with the highest frequency (i.e., the “mode”). In all other regards, we followed the same procedure as for Experiment 1.

## 4 Results

### 4.1 Experiment 1: Designing UPP2

In this first experiment we evaluated variants of UPP2, varying (a) use of adjusted or raw bit-scores, (b) using MAGUS or PASTA backbones, (c) changing the value for *z* (maximum allowed size of subsets before decomposition stops), and (d) use of EarlyStop or Hierarchical as opposed to all-against-all. On all the datasets we explored (i.e., full-length and also HF versions of 1000M1 and RNASim1000), there were no noteworthy differences in alignment accuracy for any of these modifications, with the exception that using MAGUS instead of PASTA for the backbone alignment improved accuracy (Supplementary Figures S14-S16).

We also saw that using MAGUS instead of PASTA for the backbone alignment reduced runtime (Supplementary Figure S15) and that using the new search strategies (EarlyStop or Hierarchical) improved runtime even further (Supplementary Figure S16). Specifically, UPP(PASTA)+adj+Hierarchical reduced the runtime by a large margin compared to UPP(PASTA)+adj and UPP(PASTA)+adj+EarlyStop further improved runtime compared to UPP(PASTA)+adj +Hierarchical.

The runtime improvement obtained through the use of Hierarchical or EarlyStop is not surprising, but the achievement of comparable accuracy was not guaranteed. Although we did not see a difference in accuracy between *z* = 2 compared to *z* = 10, because previous studies (e.g., Mirarab *et al*. (2012)) has suggested the potential for this setting to improve accuracy, we set *z* = 2 for the default for all datasets except the two largest Homfam datasets where we used *z* = 10 due to constraints on computational resources. Our final default settings for the algorithmic parameters are to use: (a) adjusted bit-scores, (b) MAGUS for the backbone alignment, and (c) EarlyStop for the search strategy. We denote this variant simply as “UPP2”. We use “UPP(MAGUS)+adj” to refer to UPP with adjusted bit-scores and MAGUS backbone alignments.

### 4.2 Experiment 2: UPP2 compared to Benchmark Methods

In this experiment we compared UPP2 to other alignment methods on the testing datasets. Experiment 2(a) explored results on simulated datasets with fragmentary sequences, Experiment 2(b) explored results on 16S biological datasets, and Experiment 2(c) explored results on Homfam biological datasets.

#### Experiment 2a: Results on simulated datasets with fragmentation

In this experiment we evaluated UPP2 to other alignment methods on the ROSE simulated datasets with fragmentary sequences; Figure 3 shows results for all methods on a representative sample of 6 model conditions, and Figure 4 shows results just for the three best methods on all 14 model conditions. On these datasets, UPP2 and UPP(MAGUS)+adj were the most accurate, followed by MAGUS. PASTA and MAFFT had comparable accuracy to each other and trailed behind the leading group of three methods. Clustal Omega, T-COFFEE, and MUSCLE were the least accurate methods, but MUSCLE was somewhat more accurate than the others, and Clustal Omega and T-COFFEE tended to perform similar to each other. There is a slight runtime advantage of using UPP2 over UPP(MAGUS)+adj on these datasets (Supplementary Figure S17). Figure 4 compares the top three methods, UPP2, MAGUS, and PASTA, on all 14 ROSE model conditions. We use an ordering on the model conditions from Liu *et al*. (2009) so that alignment error rates generally increase from left-to-right. While the three methods have nearly perfect alignment error and are close to identical on the six easiest model conditions (i.e., the leftmost conditions), as we move from left-to-right we see error rates increasing for all methods. However, UPP2 error rates increase more slowly than for the others, and UPP2 has the best accuracy of these three. Thus, across all the more difficult model conditions, we see that UPP2 is much more accurate than MAGUS, which in turn is more accurate than PASTA. Furthermore, the difference in accuracy between UPP2 and the next best method is often very large. In addition, while there are conditions where MAGUS is statistically significantly more accurate than UPP2, those conditions are also ones where alignment error rates are below 0.1%.

**Figure 3:**
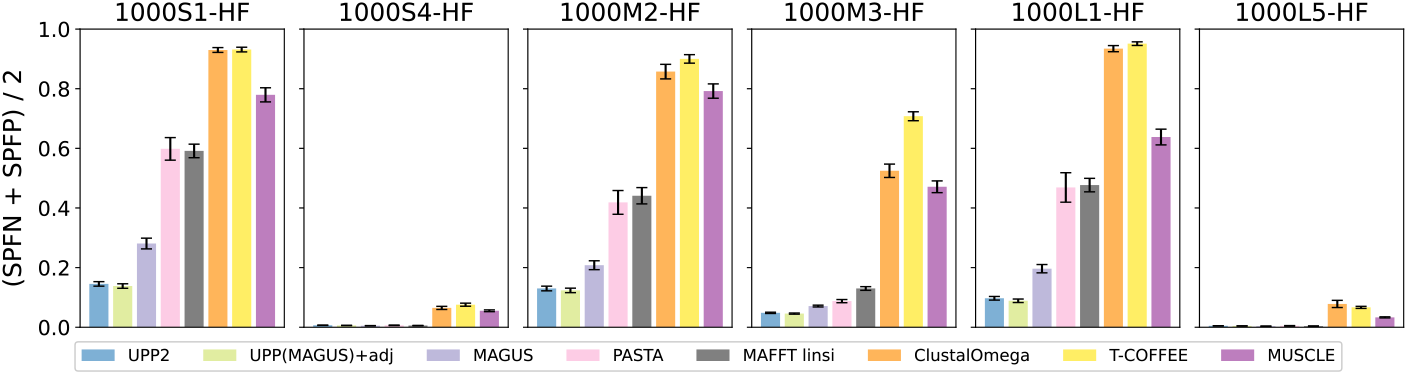
Experiment 2a: Comparison of UPP2 and Benchmark Methods on Simulated Fragmentary Datasets. All methods except T-COFFEE and MUSCLE were run in their default modes and with 16 threads, when possible. T-COFFEE was run using the default regressive mode and MUSCLE was limited to two iterations. UPP(MAGUS)+adj and UPP2 both use MAGUS backbone alignments, FastTree backbone trees, and adjusted bit-scores, but they differ in their search strategies (EarlyStop or all-against-all). All datasets have 20 replicates each. The means are shown with error bars indicating standard error for alignment error.

**Figure 4:**
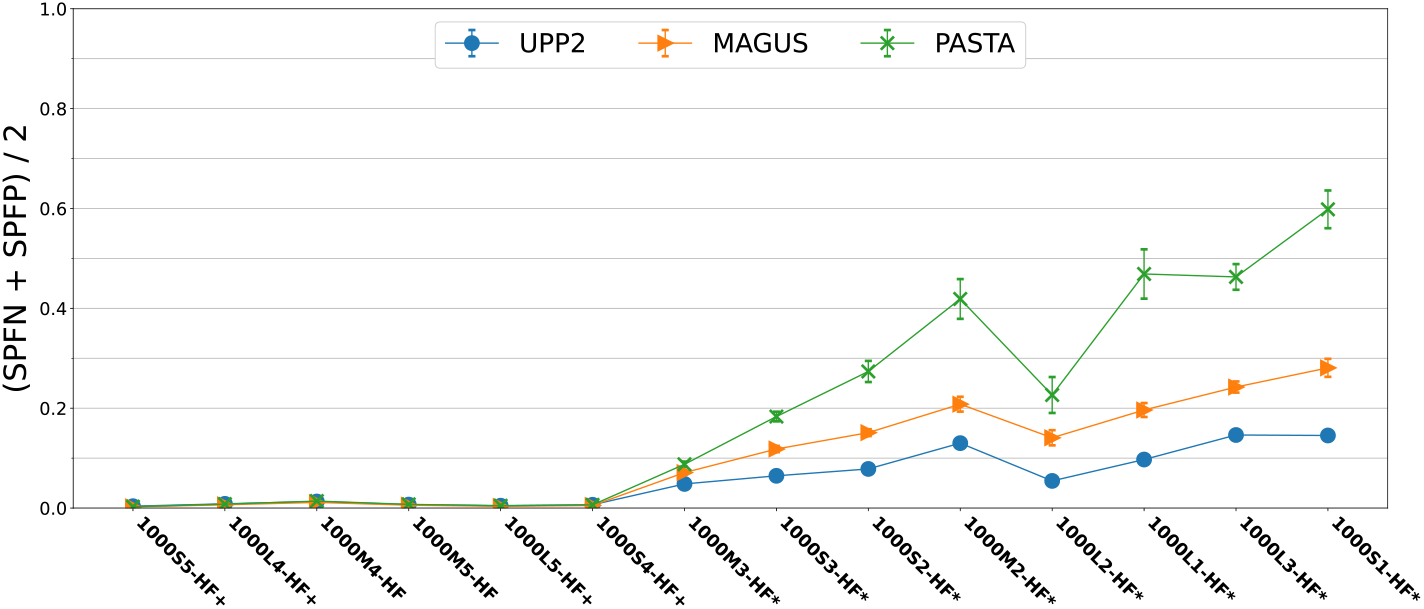
Experiment 2a: Alignment error (means across 20 replicates) of UPP2, MAGUS, and PASTA on simulated 1000-sequence datasets with fragmentary sequences. At *α* = 0.05, asterisks denote the model conditions on which UPP2 was statistically significantly better than MAGUS while plus symbols denote the model conditions on which MAGUS was statistically significantly better than UPP2; p-values are provided in Supplementary Materials Table S3. The error bars indicate standard error.

An examination of the properties of these model conditions (Supplementary Materials Table S1) shows that these conditions vary significantly in terms of average and maximum p-distances (i.e., normalized Hamming distances), and that these average p-distances generally increase as we move from left to right. Specifically, the first six model conditions (i.e., 1000S4, 1000S5, 1000M4, 1000M5, 1000L4, and 1000L5) range in average p-distances from 49.5% to 50.1%, and the next eight model conditions have increasing average p-distances that range from 66.0% to 69.6%. Thus, increases in average p-distance result in increases in alignment error for all methods, and also increase the gap between methods.

#### Experiment 2b: Results on 16S biological datasets

Results on the 16S biological datasets (Figure 5) show that UPP2 and MAGUS were the most accurate methods, with PASTA being as accurate as the top methods on 16S.B.ALL and 16S.3 but not on 16S.T. UPP(MAGUS)+adj was as accurate as UPP2 on all three 16S datasets but took far more time compared to UPP2 (about 142 hours compared to 9 hours) on the 16S.B.ALL dataset. Clustal Omega had the highest alignment error across all 16S datasets. T-COFFEE, MUSCLE, and MAFFT all performed similarly to each other on 16S.3, but MAFFT was able to beat the other two methods on 16S.B.ALL and 16S.T. Although UPP(MAGUS)+adj tied for most accurate when it could complete, it was vastly slower on the largest 16S biological dataset. UPP2 and MAGUS reliably had good accuracy (tying for best) and completed within reasonable times. The comparison between UPP2 and MAGUS shows indistinguishable accuracy on these datasets.

**Figure 5:**
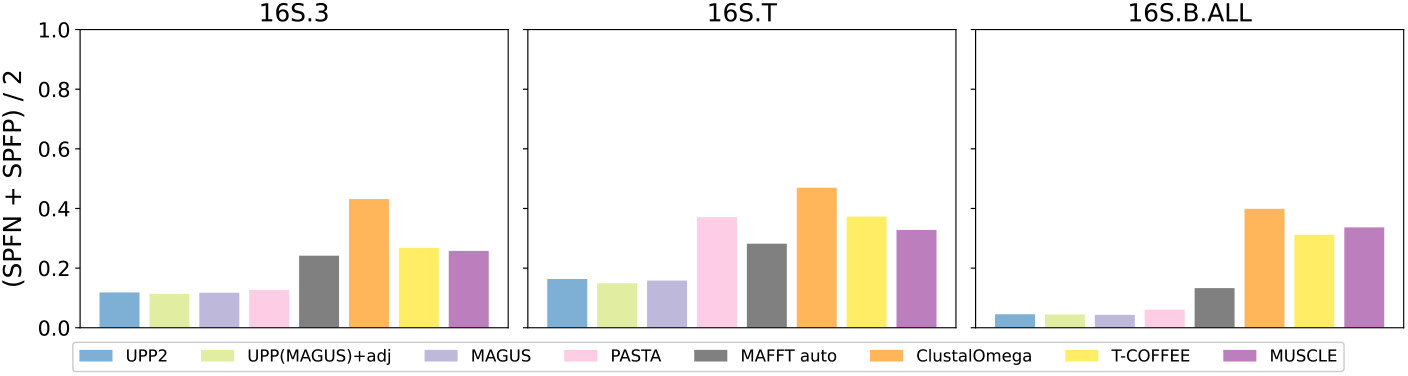
Experiment 2b: Comparison of UPP2 to other MSA Methods on 16S Biological Datasets. The three datasets are from the Comparative Ribosomal Website (Cannone *et al*., 2002). 16S.3 has 6323 sequences, 16S.T has 7350 sequences, and 16S.B.ALL has 27,643 sequences. MAFFT *auto* mode was used rather than the *linsi* mode due to the large dataset sizes.

**Figure 6:**
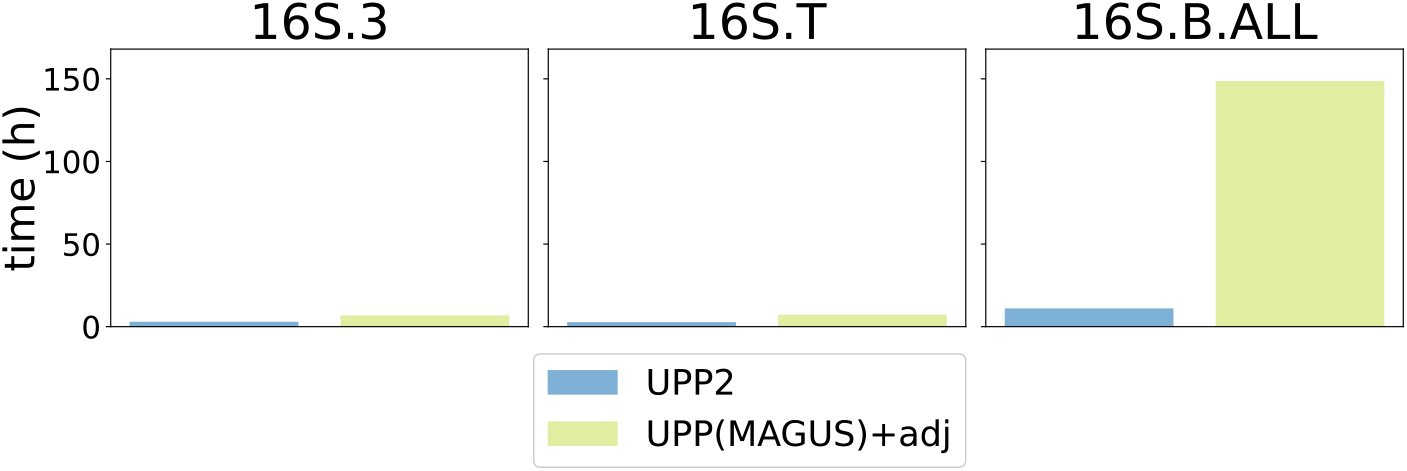
Experiment 2b: Runtime Comparison of UPP2 to UPP(MAGUS)+adj on 16S Biological Datasets. The three datasets are from the Comparative Ribosomal Website (Cannone *et al*., 2002). 1000 sequences were chosen for the backbone for 16S.3 and 16S.T while 10,000 sequences were chosen for the backbone for 16S.B.ALL. 16S.3 has 6323 sequences, 16S.T has 7350 sequences, and 16S.B.ALL has 27,643 sequences.

#### Experiment 2c: Results on Homfam biological datasets

We provide a comparison of average performance (alignment error and runtime) for methods, averaged across the ten largest Homfam datasets (Figure 7); results for individual Homfam datasets are provided in the Supplementary Materials, Figures S19-S21. This experiment includes results for both default settings for the backbone size (1000 or 10,000 sequences, depending on the number of sequences) and also for using “all” the full-length sequences in the backbone. MUSCLE failed to run on 2 of the 10 datasets (memory issues), and so results including MUSCLE are restricted to the 8 datasets on which it could run. T-COFFEE using regressive in its default mode could not complete on any of the 10 datasets, with 8 failures due to memory and the other 2 failures due to other issues. See Supplementary Materials Section S6 for full details.

**Figure 7:**
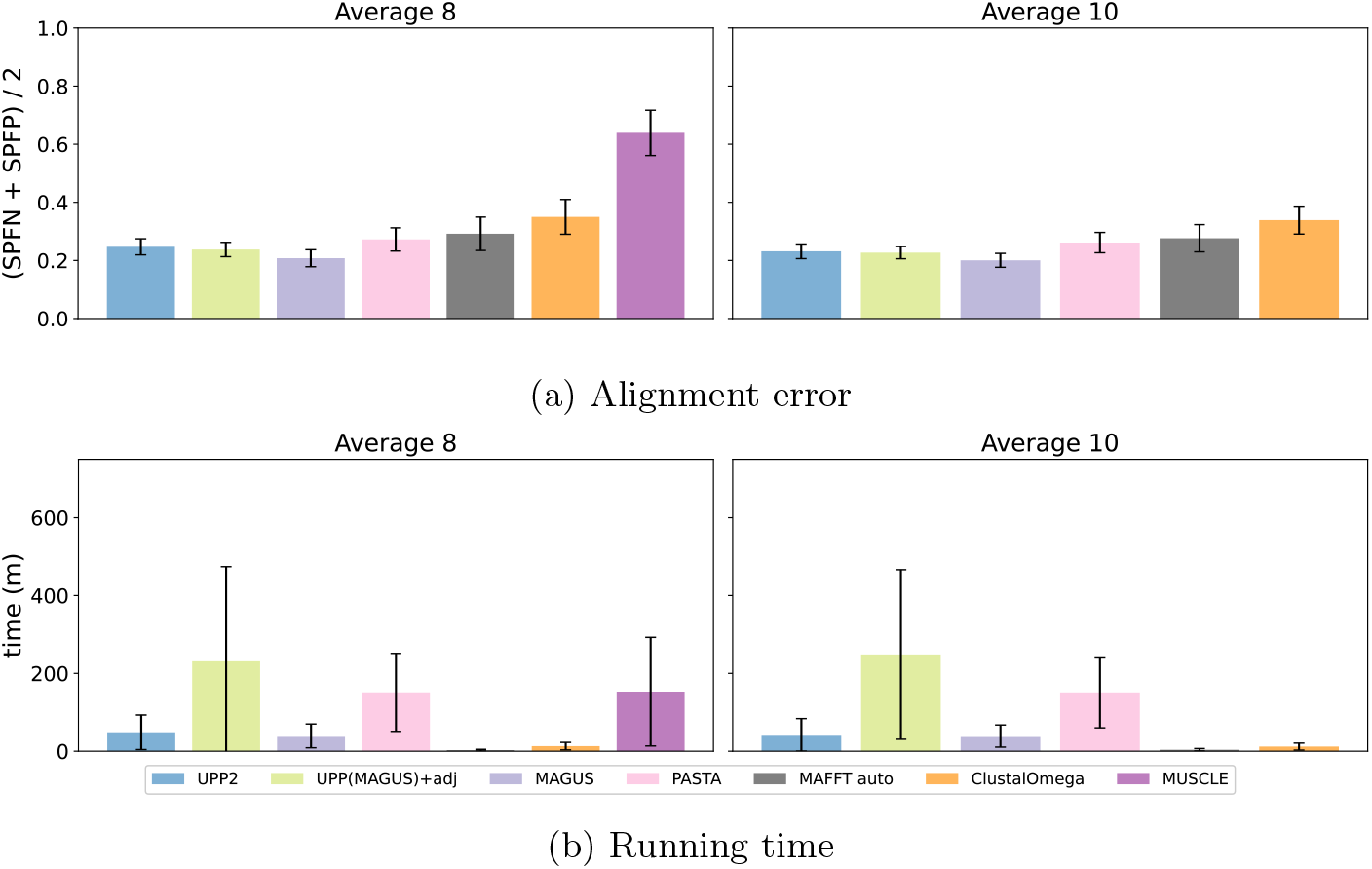
Experiment 2b: Comparison of UPP2 to other MSA Methods on Homfam Biological Datasets. Among the full-length sequences, 1000 sequences were chosen for the backbone for the four smallest datasets while 10,000 sequences were chosen for the backbone on the six larger datasets. MUSCLE could not run on the two largest datasets (zf-CCHH and rvp), but the other methods completed on all the datasets. We show average alignment error and runtime for all methods on the eight datasets where all methods completed (left) and then alignment error and runtime on all ten datasets with MUSCLE omitted (right). The number of sequences are as follows: PDZ (14,950), blmb (17,200), p450 (21,013), adh (21,331), aat (25,100), rrm (27,610), Acetyltransf (46,285), sdr (50,157), zf-CCHH (88,345), rvp (93,681). MAFFT *auto* mode was used rather than the *linsi* mode due to the large dataset sizes.

When used with the default setting for backbone size, MAGUS, UPP2, and UPP(MAGUS)+adj were the most accurate methods, with a slight advantage to MAGUS. PASTA and MAFFT–auto were close and slightly less accurate than the top three methods. Clustal-Omega was less accurate than both PASTA and MAFFT-auto, and MUSCLE had the highest error rate. MAFFT–auto was the fastest with Clustal-Omega only slightly slower, followed closely by MAGUS. UPP2 and MUSCLE followed (with about the same runtime (on those datasets on which MUSCLE completed), and then UPP(MAGUS)+adj came in last.

We also evaluated UPP2 when used with backbone alignments containing all the sequences considered full-length (instead of limited to 1000 or 10,000 sequences, depending on the dataset size). Although the impact on individual datasets varied, the use of the “ALL” setting for the backbone produced an improvement in alignment accuracy (Supplementary Materials Table S4). For example, UPP2 using default backbones had an average SPFN error across the ten datasets of 34.4% but using all full-length sequences resulted in average SPFN error of 31.5%. The use of “ALL” instead of the default setting also impacted the runtime, on average increasing the runtime by a factor of about 4 (from 42 minutes to 170 minutes). Moreover, using UPP2 with the default setting (1000 or 10,000 sequences for the backbone) allowed it to finish in at most 108 minutes on every dataset (Supplementary Materials Table S5), while using the “ALL” backbone used up to 429 minutes.

## 5 Discussion

The difference between UPP2 and UPP(MAGUS)+adj is just the replacement of the all-against-all search strategy by EarlyStop. This difference does not seem to impact accuracy in any of the model conditions or biological datasets we examined, but does allow UPP2 to have a runtime advantage over UPP(MAGUS)+adj that can be very large in some conditions. For example, on the largest 16S dataset (16S.B.ALL), UPP2 is able to complete its analysis several days before UPP(MAGUS)+adj is able to complete, and has the same alignment accuracy. The runtime advantage of UPP2 over UPP(MAGUS)+adj is also observed on the Homfam datasets, especially when using the default backbone size. However, when the backbone is large but the number of query sequences is relatively small (as is the case for some Homfam datasets when we put all the “full-length” sequences in the backbone and there were very few other sequences), then the runtime advantage, although present, is reduced.

On datasets with fragmentation, UPP2 and also UPP(MAGUS)+adj tend to have the best accuracy of all tested methods and MAGUS is very close (and sometimes better). However, the relative accuracy depends on the degree of fragmentation in the dataset as well as the rate of evolution (as reflected in the average p-distance). When there are only a small number of short sequences or when the rate of evolution is sufficiently low, then MAGUS can be as accurate as UPP2 and can even surpass UPP2 in accuracy. However, UPP2 provides an accuracy advantage over MAGUS and other standard MSA methods for those datasets exhibiting both high rates of evolution and fragmentation. The close accuracy between UPP2 and MAGUS is the result of UPP2 using MAGUS to align the backbone sequences, and is unsurprising.

This study suggests many directions for future research. At the most basic, comparisons between UPP2 and alignment methods not explored in this study, including those alignment methods that use structure information, are also needed. However, the flexible modular design of UPP2 means that variants of UPP could be explored that use advanced alignment methods, either directly for the backbone alignment or integrated into MAGUS (which is itself a flexible modular pipeline). Combining the divide-and-conquer strategies in MAGUS and UPP might enable otherwise computationally intensive methods such as the Bayesian method BAli-Phy (Suchard and Redelings, 2006) to scale to very large datasets while maintaining high accuracy (see Nute and Warnow (2016) for an early attempt of this sort). To some extent this is already enabled within UPP2, since UPP2, like its predecessors (UPP, PASTA, and SATé (Liu *et al*., 2009)) has been designed to enable the user to provide a backbone alignment. Hence, a user could select the backbone sequences and construct the alignment on those sequences, using any desired technique, and provide this backbone alignment to UPP2. Future enhancements to UPP2 could include the use of other techniques than HMMER for building the ensemble of HMMs to represent the backbone alignment (e.g., HH-suite (Steinegger *et al*., 2019) and Infernal (Nawrocki and Eddy, 2013)) and to add remaining sequences into the backbone alignment (e.g., such as the technique used in WITCH (Shen *et al*., 2022)).

## 6 Conclusions

The estimation of multiple sequence alignments on large datasets is a common step in much biological discovery. However, many modern biological datasets exhibit substantial sequence length heterogeneity, and only a few methods have been able to provide good accuracy under these conditions. The previous most accurate method for aligning datasets with fragmentary sequences was UPP, but UPP’s all-against-all approach made it computationally intensive. By replacing this search strategy with the EarlyStop approach, UPP2 achieves the same high accuracy but is much faster than UPP(MAGUS)+adj. The improvement in runtime without degradation of accuracy provided by UPP2 is encouraging, and suggests the potential for even more significant advances.

This study, taken as a whole, suggests that UPP2 is a useful method for multiple sequence alignment that generally matches or improves on the accuracy of other methods on large datasets, and that provides distinct improvements over the next best method, MAGUS, on datasets with many fragmentary sequences (especially but not only when there are high rates of evolution). UPP2 is faster than UPP due to its modified search strategy, but slower than MAGUS. Fortunately, UPP2 is scalable to large datasets, and completes within a few hours even on the largest datasets we studied with more than 90,000 sequences. Thus, the choice between MAGUS and UPP2 would need to depend on the dataset size, degree of sequence length heterogeneity (and specifically inclusion of fragmentary sequences) and rate of evolution. Given the increased availability of sequence data and the benefits obtained through dense taxon sampling, methods such as UPP2 should provide valuable functionality to many studies.

## Supporting information

Supplementary materials for UPP2

