## Supplementary materials for UPP2 for "UPP2: Fast and Accurate Alignment Estimation of Datasets with Fragmentary Sequences"

[1]Department of Computer Science, University of Illinois Urbana-Champaign, Champaign, IL 61820
*To whom correspondence should be addressed.

#### Contents

**S1 Additional Details about UPP and UPP2** **3**

**S2 Adjusted bit-scores** **4**

**S3 Method Versions and Commands** **5**

**S4 Dataset properties** **8**

**S5 Additional Results** **16**

**S6 Information about MSA failures** **33**

### S1 Additional Details about UPP and UPP2

Here we describe some additional details about UPP and UPP2 that were not provided in the main paper due to space limitations.

UPP2 uses the same *hmmsearch* parameters as UPP, such as setting an extremely high e-value cut-off and turning off all filters, in order to increase the probability of assigning each query sequence to an HMM. If a query sequence has no bit-score for a given HMM, then the bit-score will default to zero. If there are ties after computing the adjusted bit-scores, then the tie will be broken randomly, thus ensuring that every query sequence is assigned to a unique HMM. Then, each query sequence is added into the subset alignment for the selected HMM using *hmmalign*. By default, *hmmalign* outputs the outer edge unaligned residues as lower case characters, which are present in the output alignment of UPP and UPP2 as well. Following UPP's procedures, unaligned residues from different sequences are collapsed into a single column in order to save space.

#### S2 Adjusted bit-scores

One of the differences between UPP2 and the earlier version (available in the original github page) is the use of adjusted bit-scores instead of "raw" bit-scores; this modification is published in [2], and provided here to be self-contained.

The bit-score of a query sequence given a HMMER HMM is $\log_2 \frac{P(q|H)}{P(q|H_0)}$ where $H$ is the HMM, $q$ is the query sequence, and $H_0$ is the null model, or the random model. Using Bayes' theorem, we arrive at the probability of $H_i$ generating sequence $q$ as follows.

$$
\begin{aligned}
P(H_i|q) &= \frac{P(q|H_i) \cdot P(H_i)}{P(q)} \\
&= \frac{P(q|H_i) \cdot P(H_i)}{\Sigma_{j=1}^{n} P(q|H_j) \cdot P(H_j)}
\end{aligned}
$$

where $n$ is the number of HMMs ($H_i ... H_n$).

If we assume that the more sequences the HMM is trained on the more likely the HMM is to output a sequence, then we can transform the above into the following.

$$
\begin{aligned}
P(H_i|q) &= \frac{P(q|H_i) \cdot \frac{s_i}{S}}{\Sigma_{j=1}^{n} P(q|H_j) \cdot \frac{s_j}{S}} \\
&= \frac{1}{\Sigma_{j=1}^{n} \frac{P(q|H_j) \cdot s_j}{P(q|H_i) \cdot s_i}} \\
&= \frac{1}{\Sigma_{j=1}^{n} 2^{\log_2 \frac{P(q|H_j) \cdot s_j}{P(q|H_i) \cdot s_i}}}
\end{aligned}
$$

where $s_i$ is the number of sequences that HMM $H_i$ was trained on and $S$ is the total number of sequences that the HMMs were trained on.

From the definition of bit-scores, we can derive the following:

$$
\begin{aligned}
BS(H_j) - BS(H_i) &= \log_2 \frac{P(q|H_j)}{P(q|H_0)} - \log_2 \frac{P(q|H_i)}{P(q|H_0)} \\
&= \log_2 \frac{P(q|H_j)}{P(q|H_i)}
\end{aligned}
$$

So

$$
\begin{aligned}
P(H_i|q) &= \frac{1}{\Sigma_{j=1}^{n} 2^{\log_2 \frac{P(q|H_j) \cdot s_j}{P(q|H_i) \cdot s_i}}} \\
&= \frac{1}{\Sigma_{j=1}^{n} 2^{BS(H_j) - BS(H_i) + \log_2 \frac{s_j}{s_i}}}
\end{aligned}
$$

As shown in [2], adjusted bit-scores are always between 0 and 1, and can be interpreted as the probability that the given HMM generates the given query sequence. In particular, the sum, across all the HMMs in the ensemble, is always 1.

#### S3 Method Versions and Commands

##### FastSP

- Version: 1.7.1
- Availability: `https://github.com/smirarab/FastSP`
- Note: -ml and -mlr flags are omitted for MAFFT alignments since MAFFT only outputs lowercase characters
- Command:

```
java -jar FastSP.jar -ml -mlr -r <REFERENCE ALIGNMENT> -e <ESTIMATED ALIGNMENT>
```

##### MAFFT

- Version: 7.487
- Availability: `https://mafft.cbrc.jp/alignment/software/`
- Command:

```
linsi --thread 16 <SEQUENCE FILE> 1> <OUTPUT>/mafft.fasta
```

##### MUSCLE

- Version: 3.8.31
- Availability: `https://www.drive5.com/muscle/`
- Command:

```
muscle -in <SEQUENCE FILE> -out <OUTPUT>/muscle.fasta
```

##### ClustalOmega

- Version: 1.2.4
- Availability: `http://www.clustal.org/omega/`
- Command:

```
clustalo --threads=16 --in <SEQUENCE FILE> --out <OUTPUT>/clustalo.fasta
```

##### MAGUS

- Commit id: 95522ec9539575189a0a2f90baaf81cbde480034
- Availability: `https://github.com/vlasmirnov/MAGUS`
- Command:

```
python MAGUS/magus.py -d <OUTPUT> -i <SEQUENCE FILE> -o <OUTPUT>/magus.fasta
```

##### T-COFFEE

- Version: Version 13.45.0.4846264
- Availability: `http://www.tcoffee.org/`
- Command:

```
t_coffee -thread=16 -reg -seq <SEQUENCE FILE> -outfile <OUTPUT>/t_coffee.fasta
```

##### PASTA

- Version: PASTA v1.9.0
- Availability: `https://github.com/smirarab/PASTA`
- Command:

```
python run_pasta.py -i <SEQUENCE FILE> --num-cpus 16 -o <OUTPUT> --temporaries \
<OUTPUT>
```

##### UPP and UPP2

- Commit ID: 05591949d78f9e98f51c000f119c4b191f2696ef
- Availability: `https://github.com/gillichu/sepp`
- Note: By setting decomp_only and bitscore_adjust as True and leaving hier_upp and early_stop as False, we are able to replicate UPP's algorithm with adjusted bit-scores.
- Command:

```
python run_upp.py -c <CONFIG FILE>
```

##### UPP2 Config

```
[commandline]
sequence_file=<QUERY SEQUENCES>
alignment=<BACKBONE FASTA>
tree=<BACKBONE TREE>
backboneSize=<BACKBONE SIZE>
alignmentSize=<DECOMPOSITION SIZE>
molecule=<dna/rna/amino>
cpu=16
tempdir=<TEMP DIRECTORY>
outdir=<OUTPUT DIRECTORY>

[upp2]
decomp_only=True
bitscore_adjust=True
hier_upp=False
early_stop=False
```

##### UPP2-Hierarchical Config

```
[commandline]
sequence_file=<QUERY SEQUENCES>
alignment=<BACKBONE FASTA>
tree=<BACKBONE TREE>
backboneSize=<BACKBONE SIZE>
alignmentSize=<DECOMPOSITION SIZE>
molecule=<dna/rna/amino>
cpu=16
tempdir=<TEMP DIRECTORY>
outdir=<OUTPUT DIRECTORY>

[upp2]
decomp_only=True
```

```
bitscore_adjust=True
hier_upp=True
early_stop=False
```

##### UPP2 (UPP2-EarlyStop) Config

```
[commandline]
sequence_file=<QUERY SEQUENCES>
alignment=<BACKBONE FASTA>
tree=<BACKBONE TREE>
backboneSize=<BACKBONE SIZE>
alignmentSize=<DECOMPOSITION SIZE>
molecule=<dna/rna/amino>
cpu=16
tempdir=<TEMP DIRECTORY>
outdir=<OUTPUT DIRECTORY>

[upp2]
decomp_only=True
bitscore_adjust=True
hier_upp=True
early_stop=True
```

#### S4 Dataset properties

Table S1: ROSE, RNASim, and 16S dataset properties. We show the average and maximum p-distances (normalized Hamming distances) and number of sequences in each of the study datasets. The ROSE and RNASim datasets are studied in two versions: unmodified (i.e., without fragmentation) and high-fragmentation (HF, where 50% of the sequences are shortened to approximately 25% of the original median sequence length). Here we show the empirical properties of the unmodified versions of these datasets.

| Name | Sim/Bio | # Sequences | avg. p-dist. | max. p-dist. |
|---|---|---|---|---|
| 1000S1 | Sim | 1000 | 0.694 | 0.768 |
| 1000S2 | Sim | 1000 | 0.693 | 0.768 |
| 1000S3 | Sim | 1000 | 0.686 | 0.763 |
| 1000S4 | Sim | 1000 | 0.501 | 0.608 |
| 1000S5 | Sim | 1000 | 0.498 | 0.611 |
| 1000M1 | Sim | 1000 | 0.695 | 0.769 |
| 1000M2 | Sim | 1000 | 0.684 | 0.762 |
| 1000M3 | Sim | 1000 | 0.660 | 0.741 |
| 1000M4 | Sim | 1000 | 0.495 | 0.606 |
| 1000M5 | Sim | 1000 | 0.499 | 0.602 |
| 1000L1 | Sim | 1000 | 0.695 | 0.769 |
| 1000L2 | Sim | 1000 | 0.696 | 0.769 |
| 1000L3 | Sim | 1000 | 0.687 | 0.763 |
| 1000L4 | Sim | 1000 | 0.500 | 0.608 |
| 1000L5 | Sim | 1000 | 0.496 | 0.606 |
| RNASim1000 | Sim | 1000 | 0.411 | 0.609 |
| 16S.3 | Bio | 6323 | 0.315 | 0.833 |
| 16S.T | Bio | 7350 | 0.345 | 0.901 |
| 16S.B.ALL | Bio | 27643 | 0.210 | 0.769 |

Table S2: **Homfam dataset properties**. We show estimates of the average and maximum p-distances (normalized Hamming distances) and number of sequences in each of the Homfam datasets. 1000 sequences were sampled from the MAGUS estimated alignments of each dataset to compute the p-distances

| Name | Sim/Bio | # Sequences | avg. p-dist. | max. p-dist. |
|---|---|---|---|---|
| PDZ | Bio | 14,950 | 0.755 | 1.0 |
| blmb | Bio | 17,200 | 0.802 | 1.0 |
| p450 | Bio | 21,013 | 0.791 | 1.0 |
| adh | Bio | 21,331 | 0.779 | 1.0 |
| aat | Bio | 25,100 | 0.810 | 1.0 |
| rrm | Bio | 27,610 | 0.763 | 1.0 |
| Acetyltransf | Bio | 46,285 | 0.801 | 1.0 |
| sdr | Bio | 50,157 | 0.747 | 1.0 |
| zf-CCHH | Bio | 88,345 | 0.635 | 0.882 |
| rvp | Bio | 93,681 | 0.140 | 0.805 |

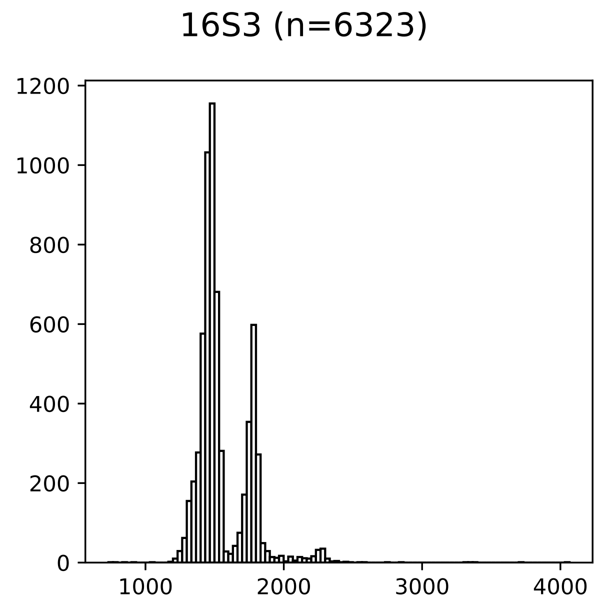

Figure S1: **Sequence length histogram for 16S.3 biological dataset** The sequence length histogram for the 16S.3 indicates at least two peaks with some sequences with length 1000 and others over 2000.

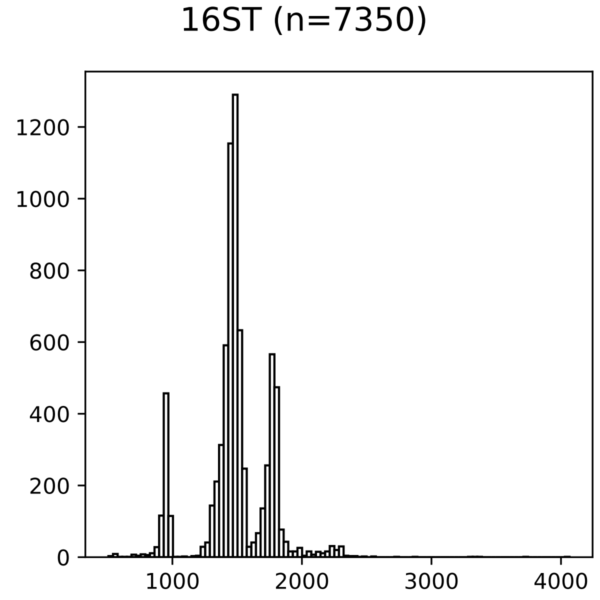

Figure S2: **Sequence length histogram for 16S.T biological dataset** The sequence length histogram for the 16S.T dataset indicates at least three peaks with one of the peaks at around 1000 and another peak closer to 2000.

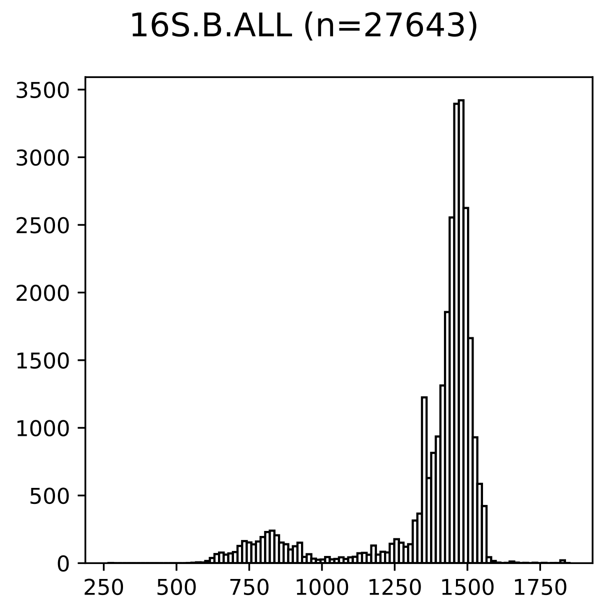

Figure S3: **Sequence length histogram for 16S.B.ALL biological dataset** The sequence length histogram for the 16S.B.ALL dataset shows moderate number of sequences with length shorter than 1250 and a peak at a little below 1500.

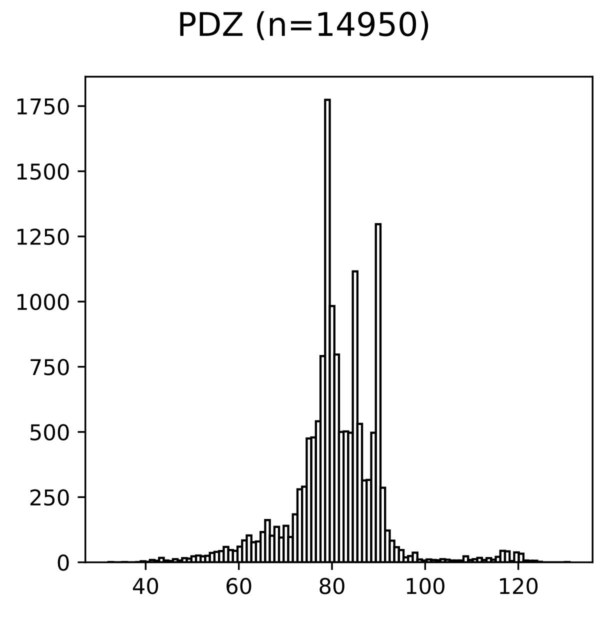

Figure S4: **Sequence length histogram for PDZ Homfam biological dataset** The sequence length histogram for the PDZ Homfam dataset shows sequences as short as 40 base pairs long as well as sequences as long as 120 base pairs long.

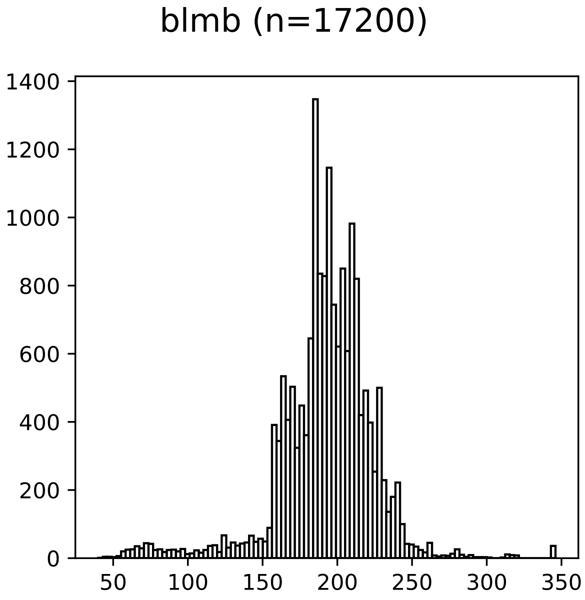

Figure S5: **Sequence length histogram for blmb Homfam Biological Dataset** The sequence length histogram for the blmb Homfam dataset shows that the majority of sequence lengths lie between 150 and 250 with some sequence lengths uniformly spread from 50 to 150.

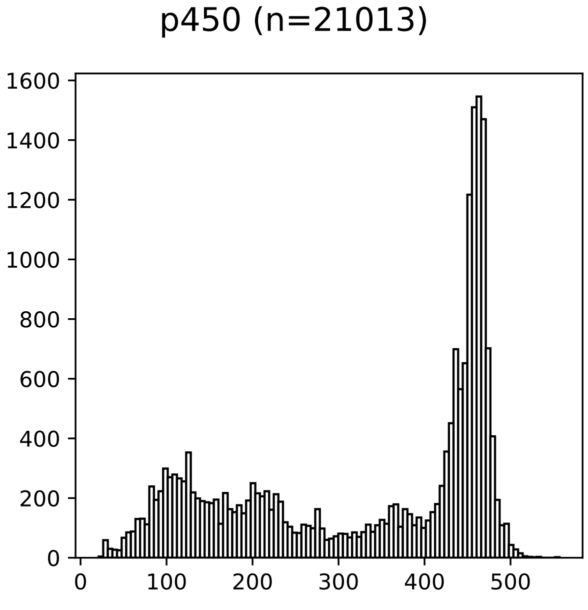

Figure S6: **Sequence length histogram for p450 Homfam Biological Dataset** The sequence length histogram for the p450 Homfam dataset shows a tall peak at around 470 with many sequences below 400 and a smaller peak at 100.

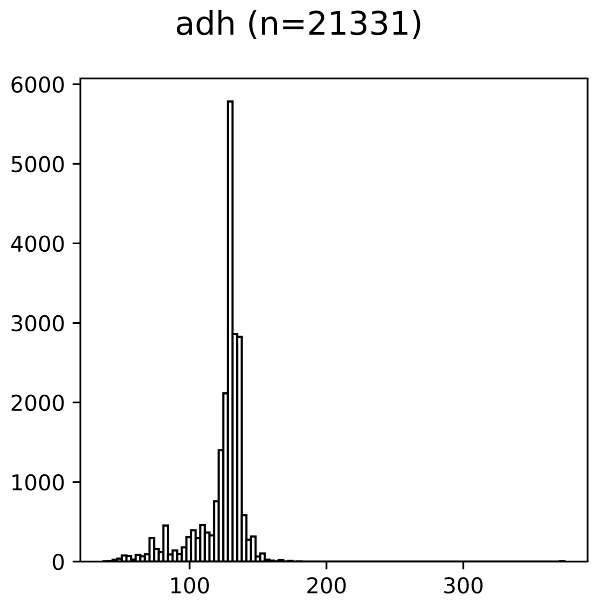

Figure S7: **Sequence length histogram for adh Homfam Biological Dataset** The sequence length histogram for the adh Homfam dataset shows a peak at around 130 with some sequences shorter than 100.

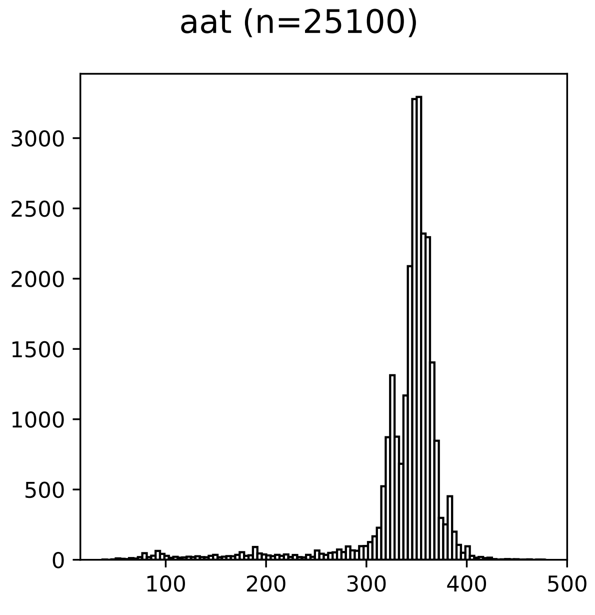

Figure S8: **Sequence length histogram for aat Homfam Biological Dataset** The sequence length histogram for the aat Homfam dataset shows a peak at around 350 with some sequences uniformly distributed from below 100 to 300.

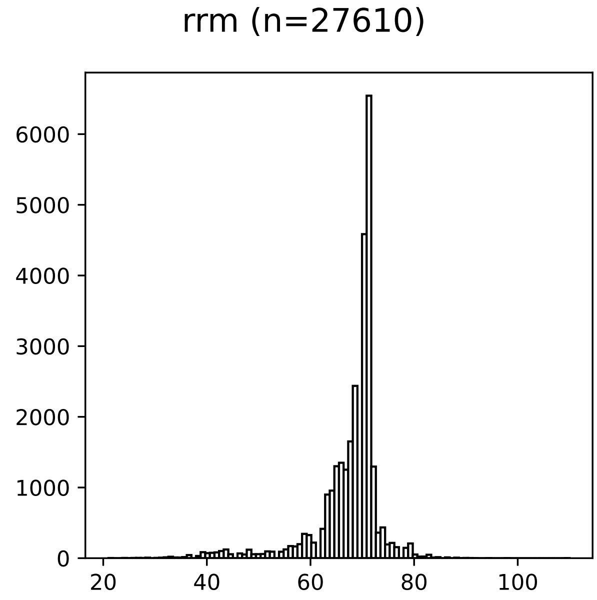

Figure S9: **Sequence length histogram for rrm Homfam Biological Dataset** The sequence length histogram for the rrm Homfam dataset shows a peak at around 70 with shorter sequences ranging from about 40 to 60.

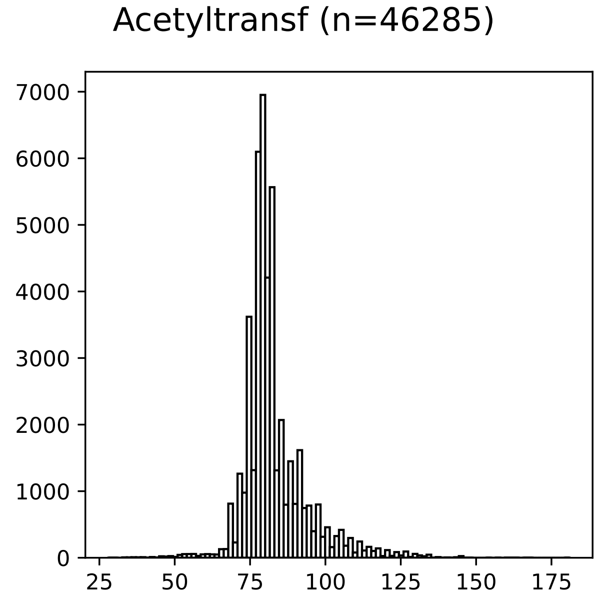

Figure S10: **Sequence length histogram for Acetyltransf Homfam Biological Dataset** The sequence length histogram for the Acetyltransf Homfam dataset shows a peak at around 80 with more sequences to the right of the peak but less so to the left of the peak.

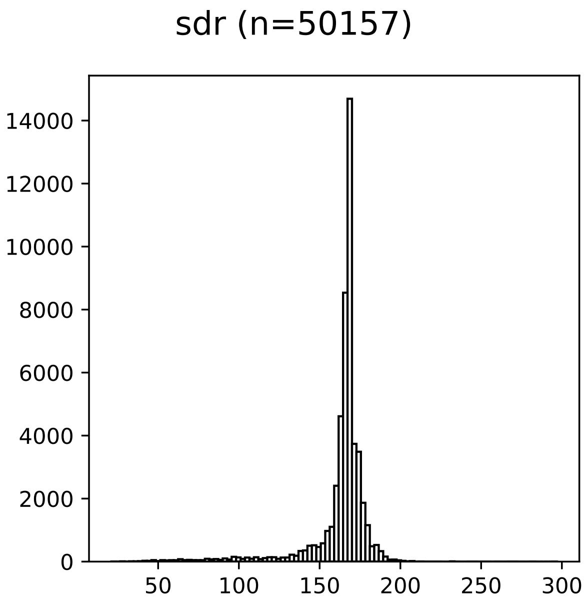

Figure S11: **Sequence length histogram for sdr Homfam Biological Dataset** The sequence length histogram for the sdr Homfam dataset shows a peak at around 170 with some sequences ranging from around 50 to 150.

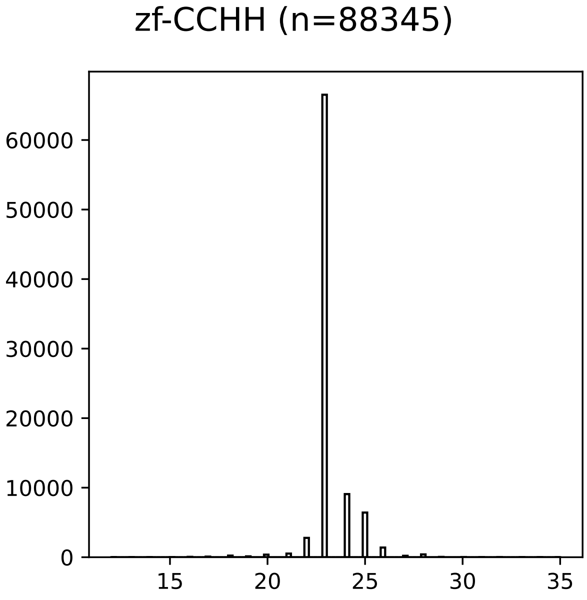

Figure S12: **Sequence length histogram for zf-CCHH Homfam Biological Dataset** The sequence length histogram for the zf-CCHH Homfam dataset shows the majority of the sequences between 20 and 25.

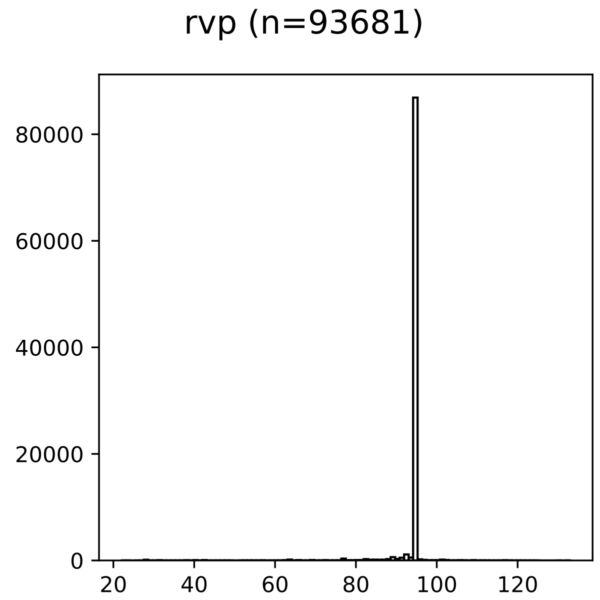

Figure S13: **Sequence length histogram for rvp Homfam biological dataset**] The sequence length histogram for the rvp dataset shows the majority of the sequences between 80 and 100 with most sequences falling under the same bin at around 95.

### S5 Additional Results

#### S5.1 Experiment 1

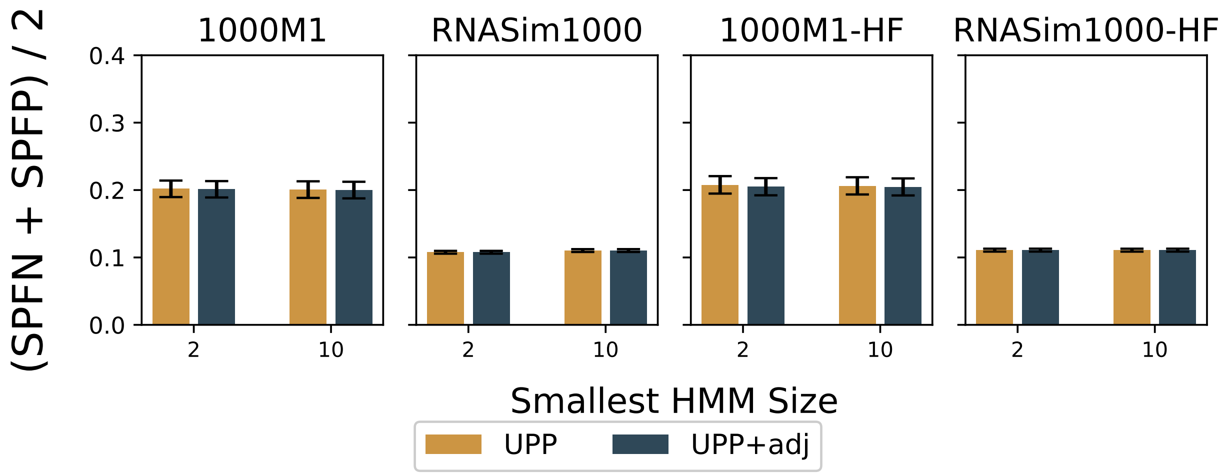

Figure S14: **Experiment 1: Impact of Adjusted Bit-score and Decomposition Size (size of smallest subsets) on Alignment Error** Both UPP and UPP+adj in this Figure uses PASTA backbone alignments. UPP uses the raw bit-scores while UPP+adj uses the adjusted bit-scores. Both methods perform an all-against-all search of HMMs to query sequences. Each subfigure shows two values for $z$, the size of the smallest subset within the decomposition strategy (i.e., decomposition size). The bar indicates the mean while the error bars indicate standard error.

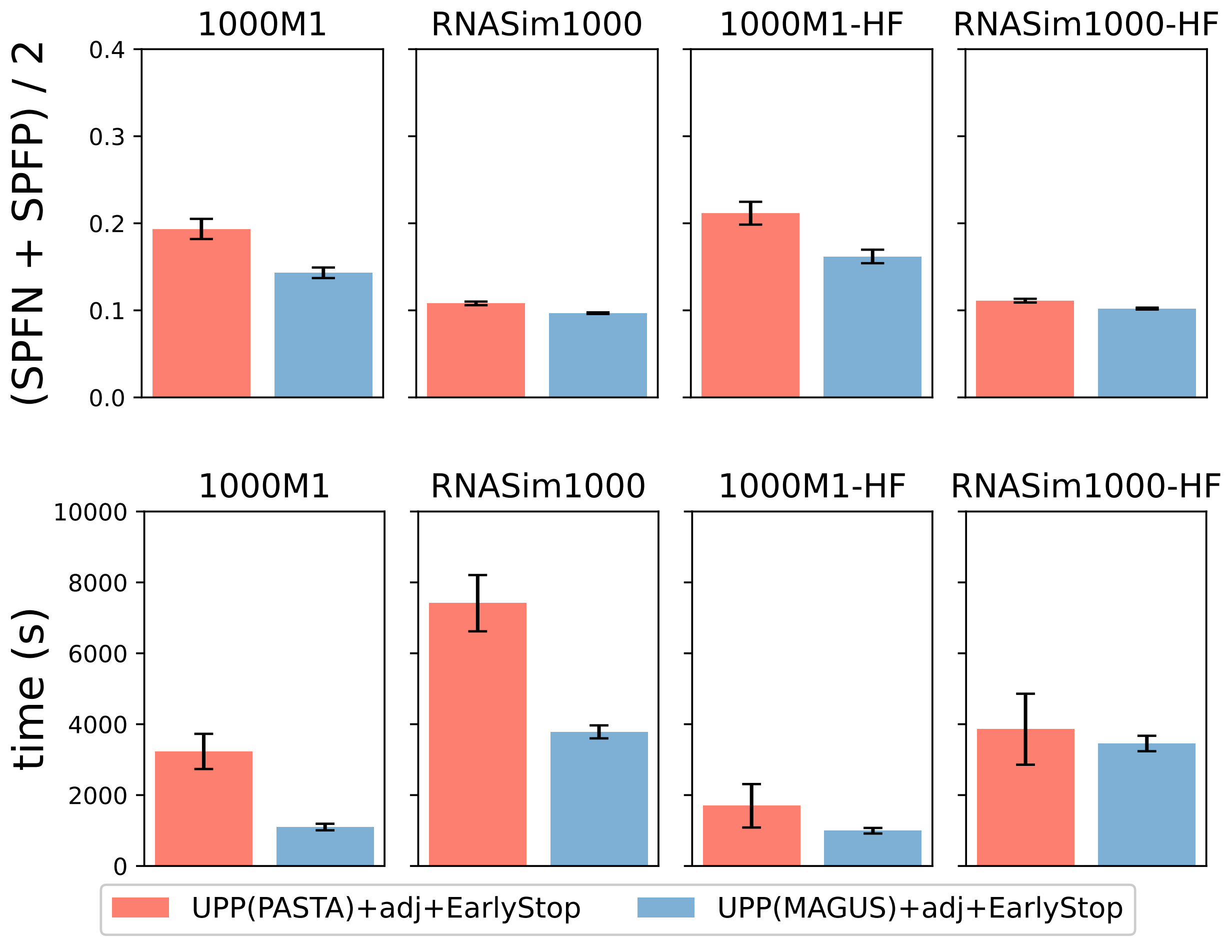

Figure S15: **Experiment 1** Impact of choice of backbone alignment method (MAGUS vs PASTA) and Decomposition Size (size of smallest subsets) on alignment accuracy and total runtime. 1000M1 has 19 replicates, RNASim1000 has 20 replicates, 1000M1-HF has 19 replicates, and RNASim1000-HF has 20 replicates. The means are shown with error bars indicating standard error for alignment error and standard deviation for running time.

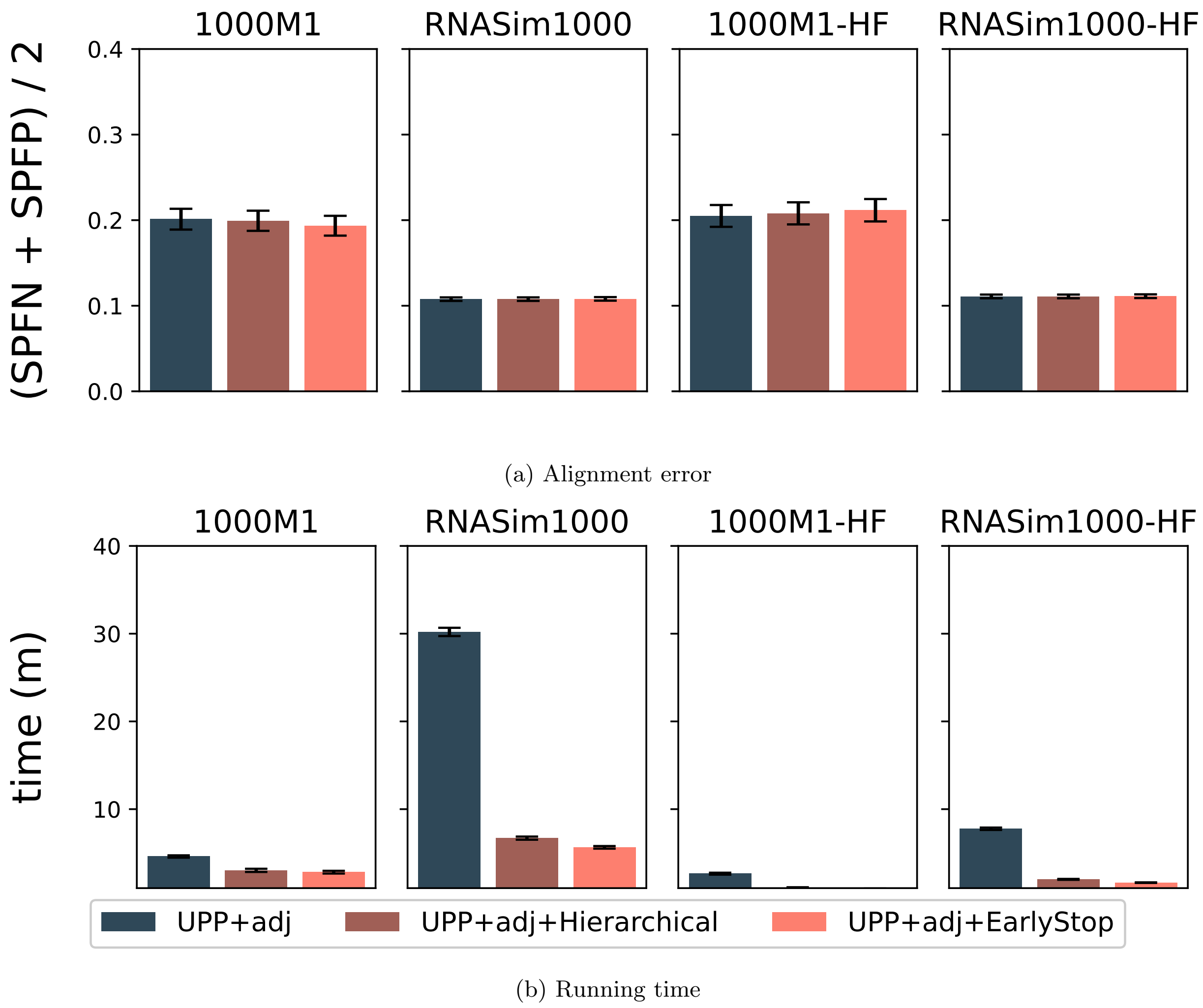

Figure S16: **Experiment 1:** Impact of Hierarchical and EarlyStop on UPP(PASTA)+adj alignment error and runtime. In this figure, UPP+adj uses the PASTA backbone alignment on full-length sequences, sets $z = 2$, and uses adjusted bit-scores. $z$ refers to the decomposition size of UPP. The runtime reported here does not include the time to compute the backbone alignment and tree. The means are shown with error bars indicating standard error for alignment error and standard deviation for running time.

#### S5.2 Experiment 2a

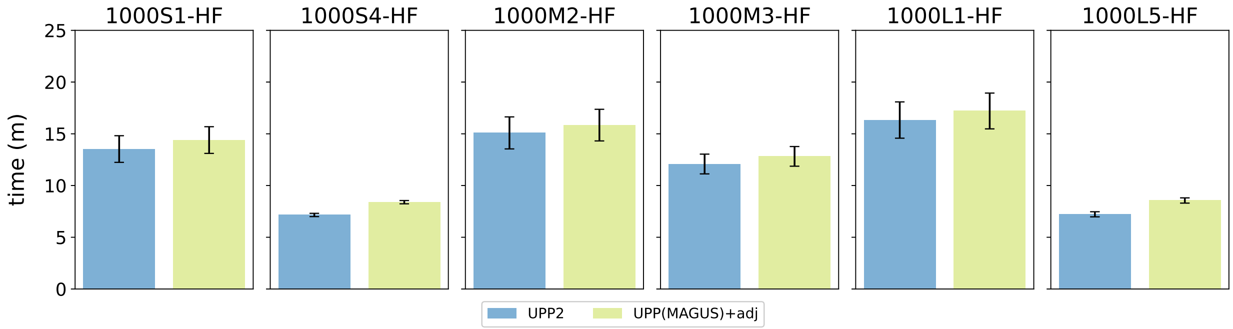

Figure S17: **Experiment 2a: Runtime comparison of UPP2 and UPP(MAGUS)+adj on simulated HF datasets** UPP(MAGUS)+adj and UPP2 both use MAGUS backbone alignments, FastTree backbone trees, and adjusted bit-scores; they differ in their search strategies (EarlyStop vs. all-against-all). All datasets have 20 replicates each. The means are shown with error bars indicating standard deviation for running time.

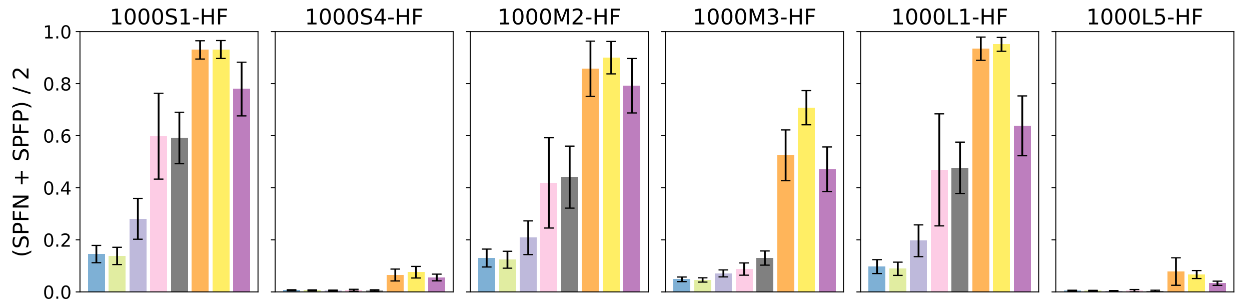

(a) Average alignment error

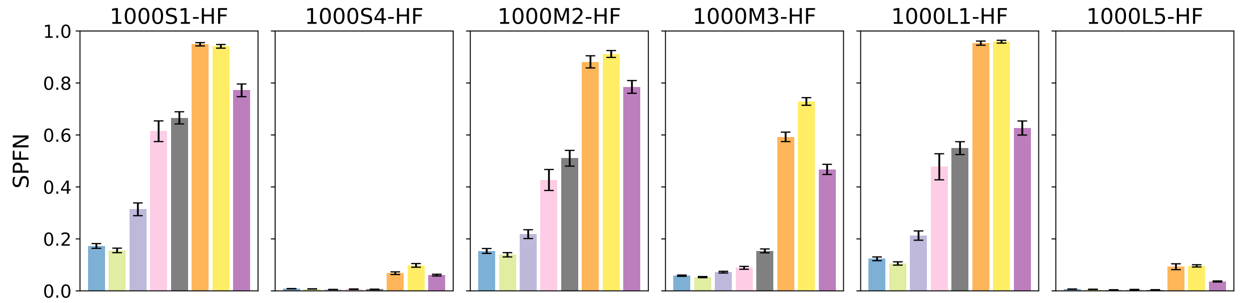

(b) SPFN alignment error

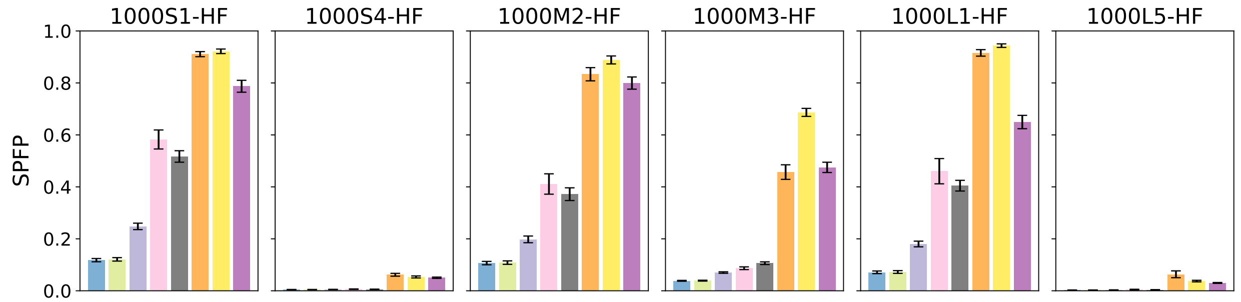

(c) SPFP alignment error

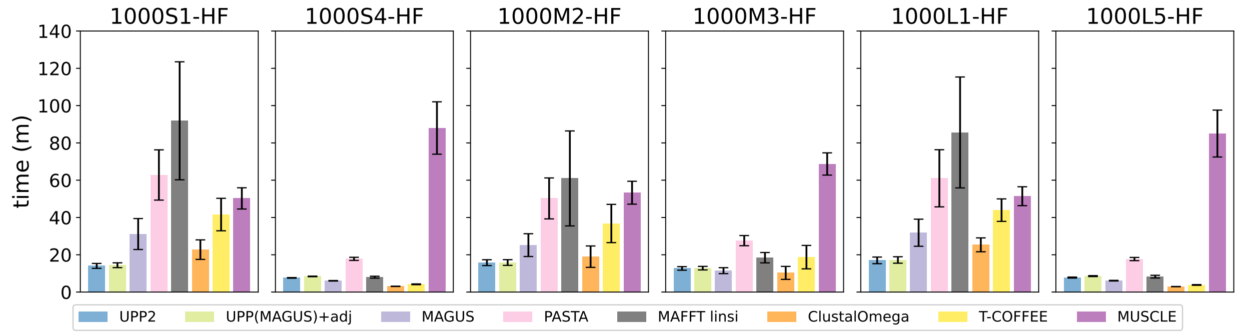

(d) running time

Figure S18: **Experiment 2: UPP2 compared to benchmark alignment methods on HF simulated datasets.** We show alignment error and runtime of UPP2 (i.e., UPP(MAGUS)+adj+EarlyStop) compared to other alignment methods. All methods except T-COFFEE and MUSCLE were run in their default modes and with 16 threads, when possible. T-COFFEE was run using the regressive mode and MUSCLE was limited to 2 iterations. All datasets have 20 replicates each. The means are shown with error bars indicating standard error for alignment error and standard deviation for running time.

Table S3: **p-values for the statistical tests in Figure 4** We show the p-values of independent two-sample t-tests of UPP2 compared to the next best method (MAGUS). A positive test statistic indicates that the next best method (MAGUS) was more accurate than UPP2 while a negative test statistic indicates that UPP2 was more accurate than the next best method (MAGUS). Note that there are many conditions where the differences are statistically significant (indicated by $p < 0.05$). However, the only cases that were statistically significant and noteworthy (i.e., differences greater than 0.01) are where UPP2 is more accurate than MAGUS.

| Name | test statistic | p-value | MAGUS error rate | UPP2 error rate |
|---|---|---|---|---|
| 1000S5-HF | 4.55 | $5.86 \times 10^{-5}$ | 0.002 | 0.004 |
| 1000L4-HF | 2.42 | $2.08 \times 10^{-2}$ | 0.007 | 0.008 |
| 1000M4-HF | 2.02 | $5.13 \times 10^{-2}$ | 0.011 | 0.013 |
| 1000M5-HF | 1.52 | $1.38 \times 10^{-1}$ | 0.006 | 0.007 |
| 1000L5-HF | 2.38 | $2.27 \times 10^{-2}$ | 0.003 | 0.005 |
| 1000S4-HF | 2.91 | $6.16 \times 10^{-3}$ | 0.005 | 0.006 |
| 1000M3-HF | -5.89 | $9.63 \times 10^{-7}$ | 0.071 | 0.049 |
| 1000S3-HF | -7.52 | $6.85 \times 10^{-9}$ | 0.118 | 0.064 |
| 1000S2-HF | -8.55 | $3.47 \times 10^{-10}$ | 0.151 | 0.078 |
| 1000M2-HF | -4.46 | $7.61 \times 10^{-5}$ | 0.204 | 0.128 |
| 1000L2-HF | -5.12 | $1.03 \times 10^{-5}$ | 0.141 | 0.054 |
| 1000L1-HF | -6.29 | $2.83 \times 10^{-7}$ | 0.198 | 0.099 |
| 1000L3-HF | -7.62 | $5.12 \times 10^{-9}$ | 0.247 | 0.149 |
| 1000S1-HF | -7.62 | $5.18 \times 10^{-9}$ | 0.271 | 0.141 |

##### S5.3 Experiment 2c

In UPP2, the backbone size is a parameter that can be controlled for the trade-off between accuracy and runtime. Since UPP2 works by aligning query sequences to the backbone alignment, the accuracy of the backbone alignment matters greatly to the final output alignment. In addition, the size of the backbone alignment, along with the decomposition size, determines how big the ensemble of HMMs is; the larger the backbone alignment and smaller the decomposition size, the bigger the ensemble of HMMs becomes.

We explore this trade-off using the Homfam datasets. Averaging across the ten datasets, using the larger backbone for both UPP2 or UPP(MAGUS)+adj improved both SPFN and SPFP compared to using the default backbone (Figure S24). On the other hand, the impact of changing the backbone size depends on the dataset (Figure S22), with some analyses improved using the larger backbone, some made less accurate, and many others not affected.

To enable a comparison to T-COFFEE Regressive, we also provide a comparison of SPFN values in Table S4, using results for T-COFFEE Regressive obtained from [1]. MAGUS provides the best accuracy, followed by UPP2 using the ALL setting, and then by MAFFT in auto mode follows. T-COFFEE Regressive and UPP2 with default settings are very close but with a small advantage to T-COFFEE. Thus, the use of the "ALL" setting for the backbone improves accuracy, making UPP2 close to the best of this set of methods, and improving on T-COFFEE Regressive and MAFFT. However, using the "default" backbone setting makes UPP2 less accurate than T-COFFEE Regresive on these 10 Homfam datasets.

The choice of backbone size impacts runtime, and the impact is different for UPP2 than for UPP(MAGUS)+adj. Increasing the backbone size to "ALL" always increased the runtime for UPP2, but had a variable impact for UPP(MAGUS)+adj: decreasing runtime for half the datasets and increasing it for the other half. Finally, UPP2 is always faster than UPP(MAGUS)+adj whether used with default backbone sizes or the "ALL" setting.

Table S4: **Homfam individual dataset SPFN error with T-COFFEE results** The SPFN error of UPP2, (All) UPP2, MAGUS, MAFFT, and T-COFFEE are shown.

| | UPP2 | (All) UPP2 | MAGUS | MAFFT | T-COFFEE |
|---|---|---|---|---|---|
| Average | 0.344 | 0.315 | 0.285 | 0.324 | 0.341 |
| PDZ | 0.265 | 0.171 | 0.214 | 0.163 | 0.320 |
| blmb | 0.354 | 0.303 | 0.298 | 0.447 | 0.776 |
| p450 | 0.404 | 0.246 | 0.263 | 0.449 | 0.359 |
| adh | 0.647 | 0.645 | 0.269 | 0.019 | 0.014 |
| aat | 0.156 | 0.180 | 0.186 | 0.274 | 0.275 |
| rrm | 0.239 | 0.234 | 0.237 | 0.292 | 0.275 |
| Acetyltransf | 0.545 | 0.549 | 0.566 | 0.551 | 0.549 |
| sdr | 0.405 | 0.457 | 0.398 | 0.544 | 0.409 |
| zf-CCHH | 0.192 | 0.184 | 0.210 | 0.213 | 0.231 |
| rvp | 0.231 | 0.177 | 0.213 | 0.288 | 0.197 |

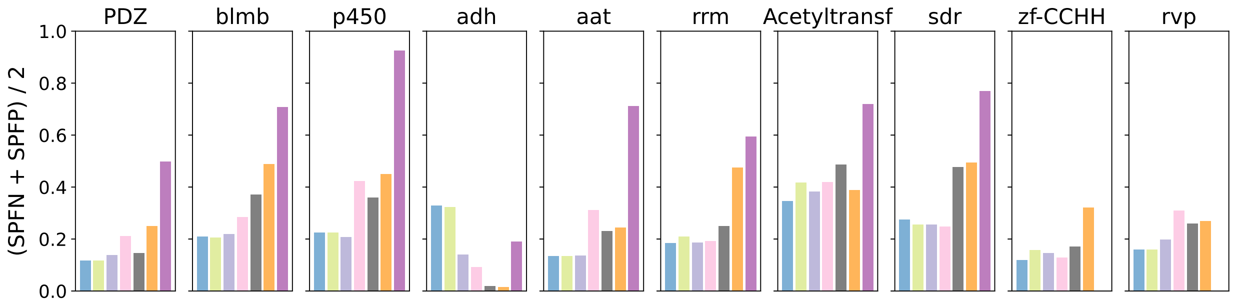

(a) Average alignment error

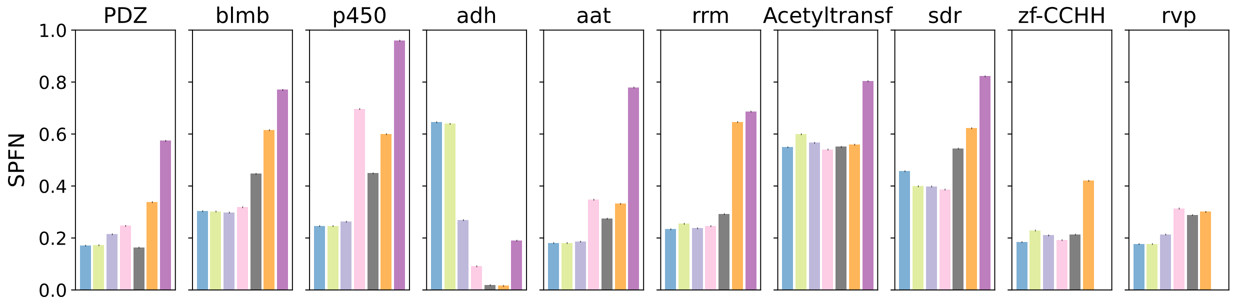

(b) SPFN alignment error

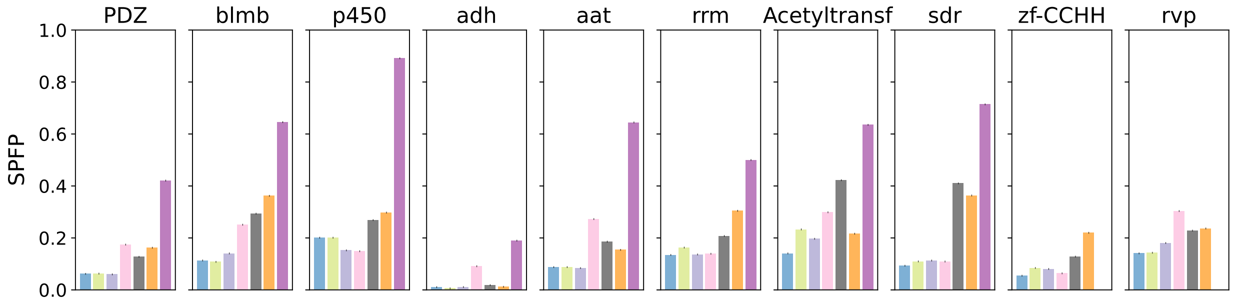

(c) SPFP alignment error

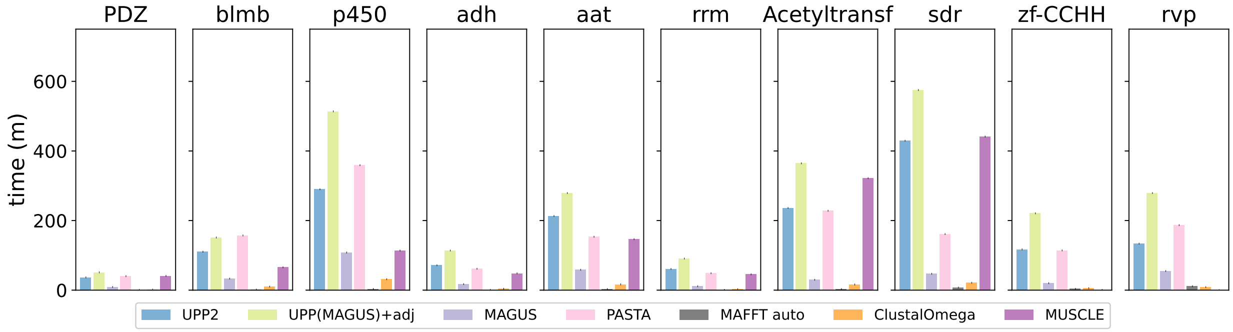

(d) Running time

Figure S19: **Alignment error and runtime of UPP variants using all full-length as backbone on the individual Homfam datasets** All full-length sequences were chosen for the backbone. UPP2 had a memory failure on zf-CCHH and rvp using a decomposition size of 2, so the results shown use a decomposition size of 10; UPP2 and UPP(MAGUS)+adj used a decomposition size of 2 otherwise. MUSCLE could not run on zf-CCHH and rvp, but the remaining benchmark methods in the legend completed on all the datasets. The number of sequences are as follows: PDZ (14,950), blmb (17,200), p450 (21,013), adh (21,331), aat (25,100), rrm (27,610), Acetyltransf (46,285), sdr (50,157), zf-CCHH (88,345), rvp (93,681).

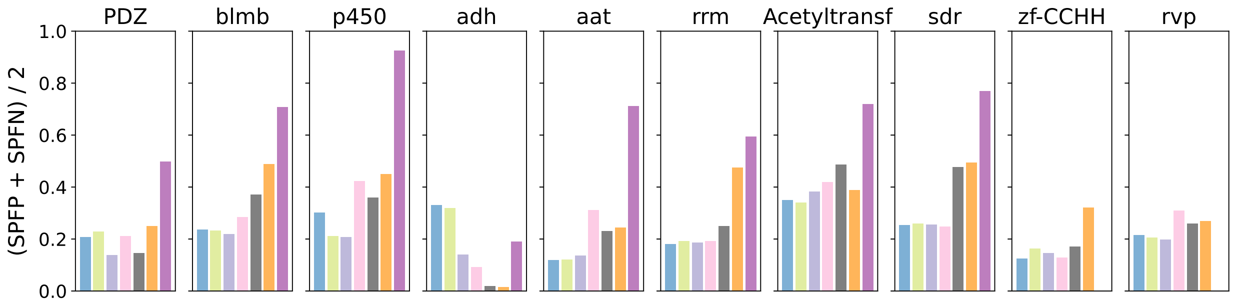

(a) Average alignment error

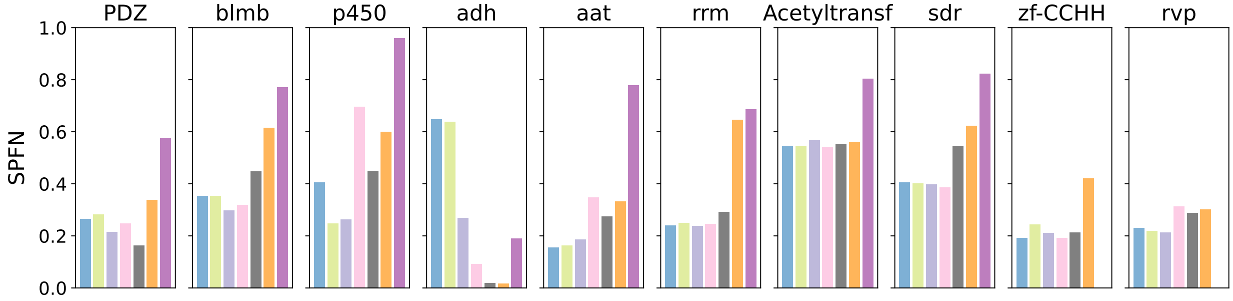

(b) SPFN alignment error

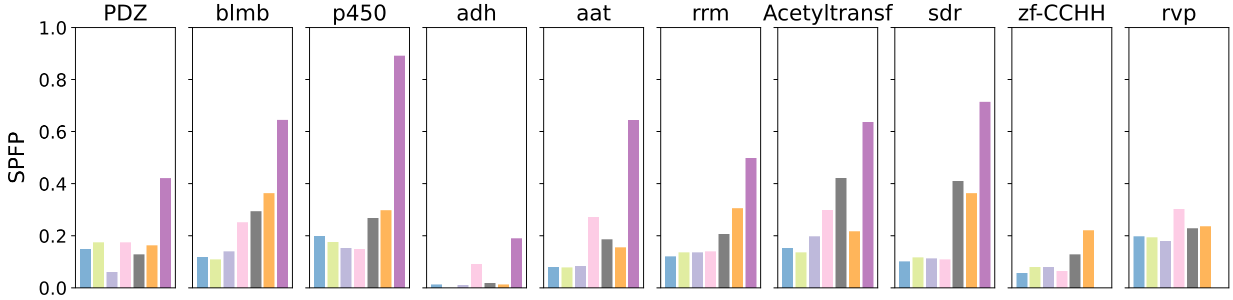

(c) SPFP alignment error

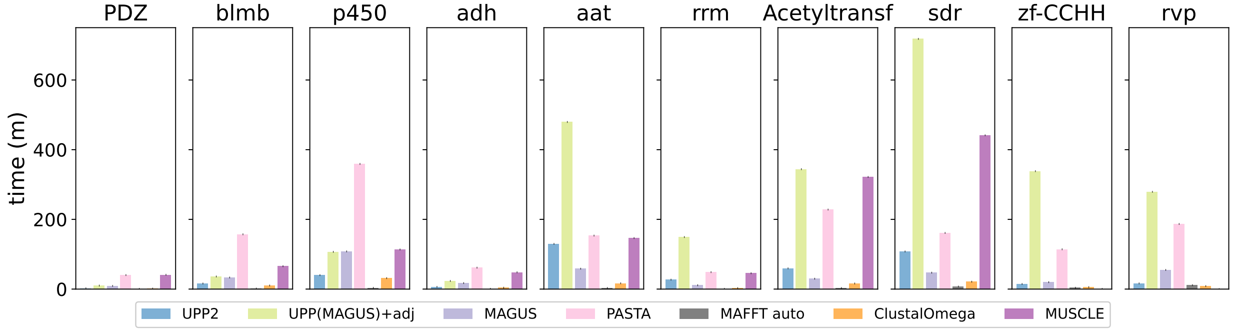

(d) Running time

Figure S20: **Alignment error and runtime of UPP variants with 1000 or 10,000 full-length as backbone on the individual Homfam datasets** Of the full-length sequences, 1000 sequences for the smallest four datasets and 10,000 sequences for the six largest datasets were chosen for the backbone. UPP2 and UPP(MAGUS)+adj used a decomposition size of 2 in all datasets. MUSCLE could not run on zf-CCHH and rvp, but the remaining benchmark methods in the legend completed on all the datasets. The number of sequences are as follows: PDZ (14,950), blmb (17,200), p450 (21,013), adh (21,331), aat (25,100), rrm (27,610), Acetyltransf (46,285), sdr (50,157), zf-CCHH (88,345), rvp (93,681).

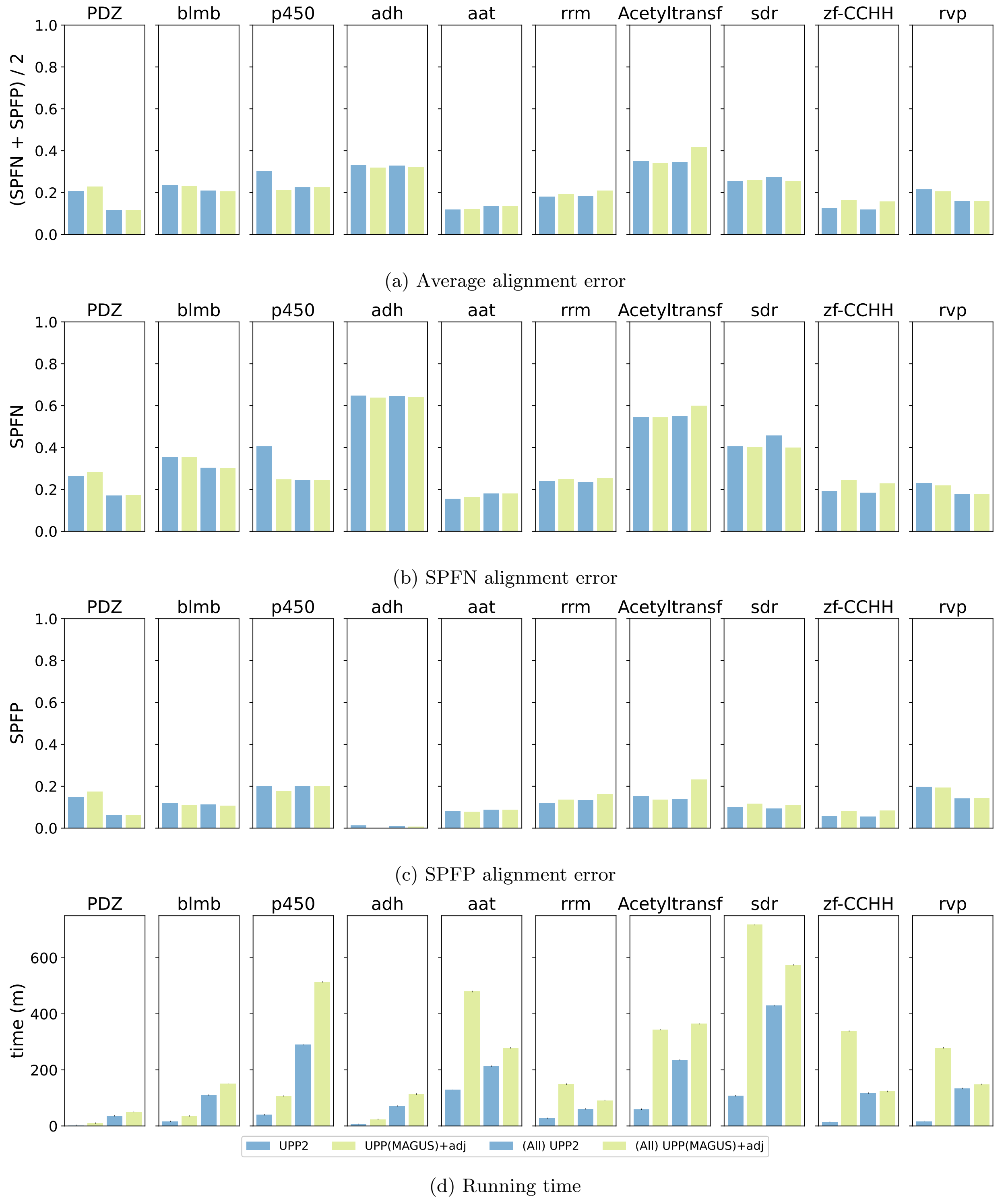

(a) Average alignment error

(b) SPFN alignment error

(c) SPFP alignment error

(d) Running time

Figure S21: **Impact of UPP2 backbone size on alignment error and runtime on the individual Homfam datasets** Default settings for the backbone (1000 or 10,000 sequences) compared to using all the “full-length” sequences. UPP2, UPP(MAGUS)+adj, (All) UPP2, and (All) UPP(MAGUS)+adj all used a decomposition size of 2 except for (All) UPP(MAGUS)+adj on zf-CCHH and rvp where the decomposition size was 10. (All) UPP(MAGUS)+adj encountered memory issues using a decomposition of size 2 on zf-CCHH and rvp. The number of sequences are as follows: PDZ (14,950), blmb (17,200), p450 (21,013), adh (21,331), aat (25,100), rrm (27,610), Acetyltransf (46,285), sdr (50,157), zf-CCHH (88,345), rvp (93,681).

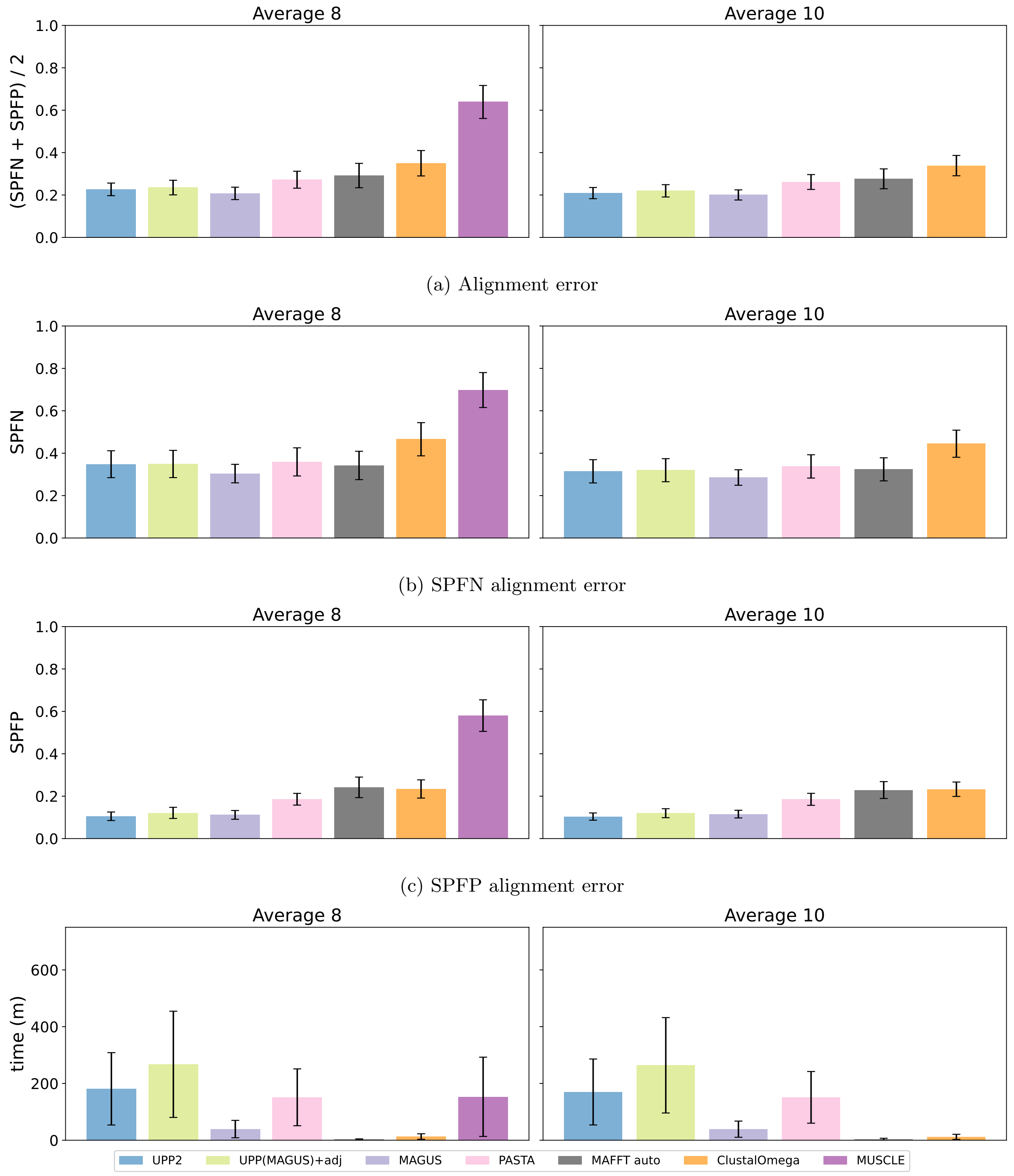

(a) Alignment error

(b) SPFN alignment error

(c) SPFP alignment error

(d) Running time

Figure S22: **Average alignment error and rntime of UPP variants with all full-length as backbone on the Homfam datasets** All full-length sequences were chosen for the backbone. UPP2 had a memory failure on zf-CCHH and rvp using a decomposition size of 2, so the results shown use a decomposition size of 10. UPP2 and UPP(MAGUS)+adj used a decomposition size of 2 otherwise. MUSCLE could not run on zf-CCHH and rvp, but the remaining benchmark methods in the legend completed on all the datasets. The results on eight datasets, excluding the two datasets which MUSCLE could not run on, were averaged together under Average 8. Excluding MUSCLE, all methods were averaged across all datasets under Average 10. The number of sequences are as follows: PDZ (14,950), blmb (17,200), p450 (21,013), adh (21,331), aat (25,100), rrm (27,610), Acetyltransf (46,285), sdr (50,157), zf-CCHH (88,345), rvp (93,681).

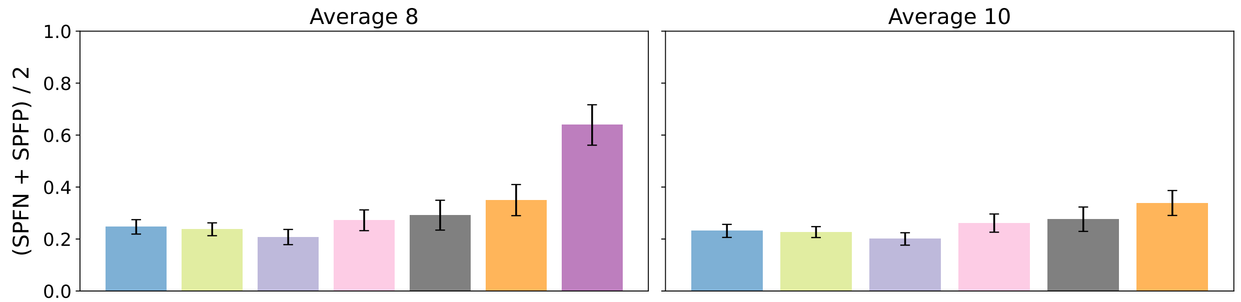

(a) Alignment error

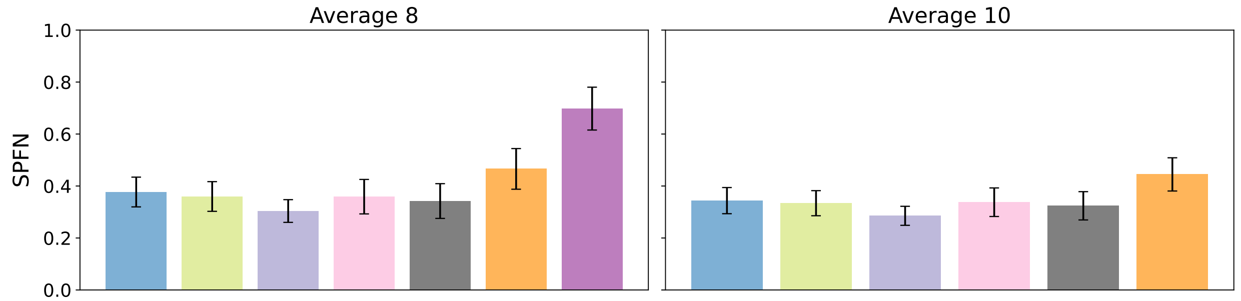

(b) SPFN alignment error

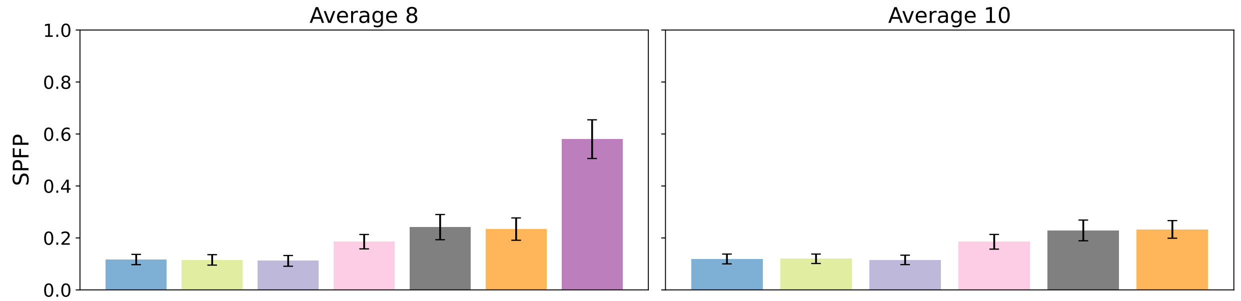

(c) SPFP alignment error

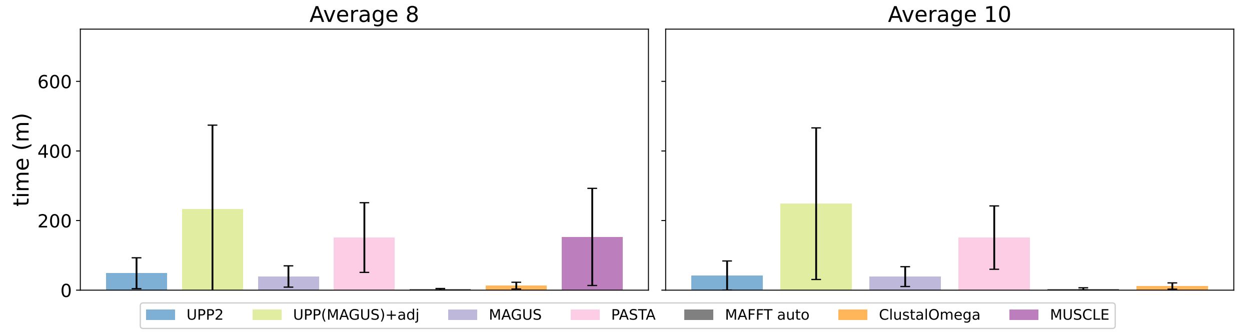

(d) Running time

Figure S23: **Average alignment error and runtime of UPP with default backbones on the Homfam datasets** From the full-length sequences, 1000 sequences were chosen for the backbone in the four smallest datasets while 10,000 sequences were chosen for the backbone. UPP2 had a memory failure on zf-CCHH and rvp using a decomposition size of 2, so the results shown use a decomposition size of 10; UPP2 and UPP(MAGUS)+adj used a decomposition size of 2 otherwise. MUSCLE could not run on zf-CCHH and rvp, but the remaining benchmark methods in the legend completed on all the datasets. The results on eight datasets, excluding the two datasets which MUSCLE could not run on, were averaged together under "Average 8". Excluding MUSCLE, all methods were averaged across all datasets under "Average 10". The number of sequences are as follows: PDZ (14,950), blmb (17,200), p450 (21,013), adh (21,331), aat (25,100), rrm (27,610), Acetyltransf (46,285), sdr (50,157), zf-CCHH (88,345), rvp (93,681).

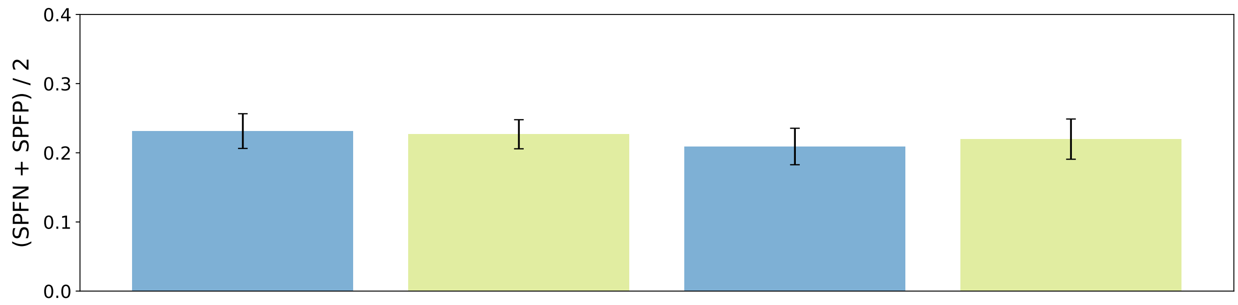

(a) Average alignment error

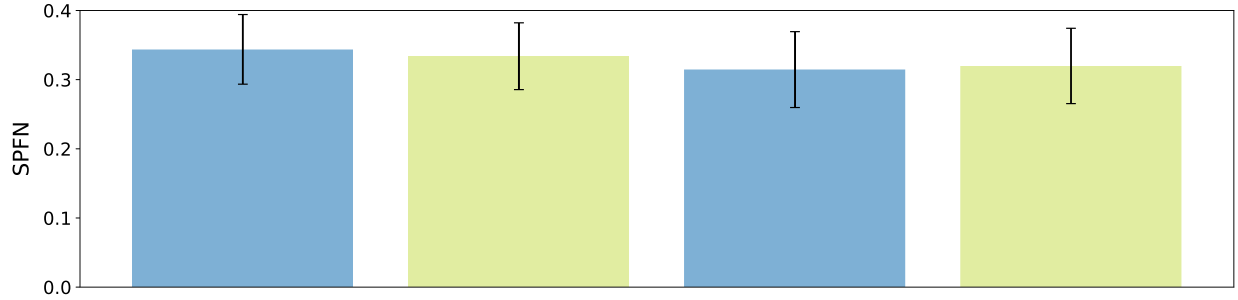

(b) SPFN alignment error

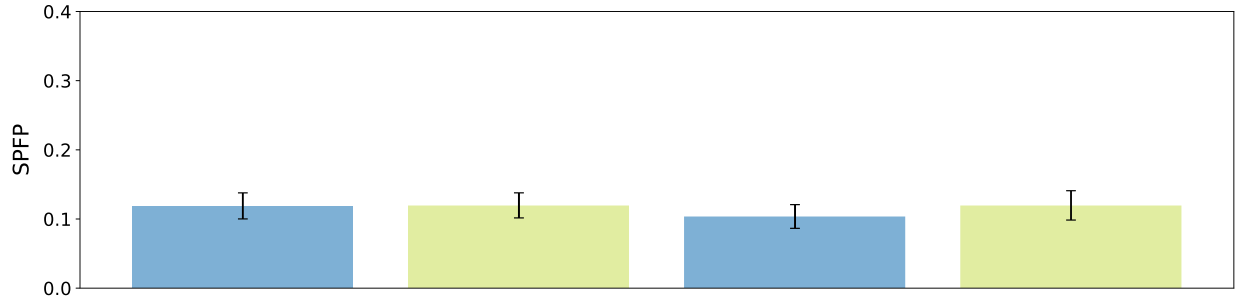

(c) SPFP alignment error

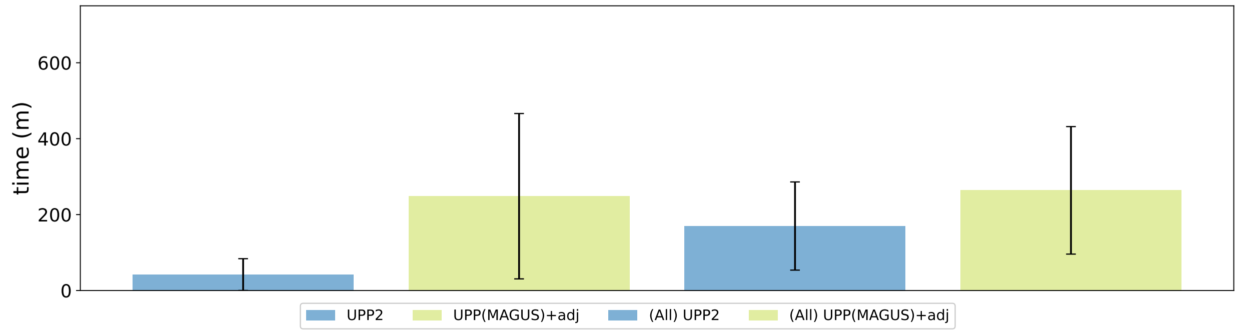

(d) Running time

Figure S24: **Impact of UPP2 backbone size on average alignment error and runtime on the Homfam datasets** All methods are averaged across the 10 Homfam datasets. For UPP2 and UPP(MAGUS)+adj, of the full-length sequences, 1000 sequences for the smallest four datasets and 10,000 sequences for the six largest datasets were chosen for the backbone. For (All) UPP2 and (All) UPP(MAGUS)+adj, all full-length sequences were chosen for the backbone. UPP2, UPP(MAGUS)+adj, (All) UPP2, and (All) UPP(MAGUS)+adj all used a decomposition size of 2 except for (All) UPP(MAGUS)+adj on zf-CCHH and rvp where the decomposition size was 10. (All) UPP(MAGUS)+adj encountered memory issues using a decomposition of size 2 on zf-CCHH and rvp. The number of sequences are as follows: PDZ (14,950), blmb (17,200), p450 (21,013), adh (21,331), aat (25,100), rrm (27,610), Acetyltransf (46,285), sdr (50,157), zf-CCHH (88,345), rvp (93,681).

Table S5: **Homfam individual dataset runtimes** The runtime of UPP2, (All) UPP2, MAGUS, and MAFFT (in minutes) are shown. Results for T-Coffee are not shown since we were unable to run T-COFFEE, and runtimes reported on other computational environments for T-COFFEE cannot be compared to runtimes reported here.

| | UPP2 | (All) UPP2 | MAGUS | MAFFT |
|---|---|---|---|---|
| Average | 42.1 | 169.9 | 39.0 | 3.7 |
| PDZ | 2.4 | 36.7 | 9.2 | 0.6 |
| blmb | 16.1 | 110.5 | 33.3 | 1.6 |
| p450 | 40.1 | 290.0 | 108.2 | 3.7 |
| adh | 6.3 | 71.4 | 17.3 | 0.8 |
| aat | 129.8 | 213.1 | 58.6 | 3.2 |
| rrm | 27.1 | 61.2 | 11.2 | 0.8 |
| Acetyltransf | 59.6 | 236.2 | 29.8 | 3.1 |
| sdr | 108.0 | 429.3 | 47.1 | 7.4 |
| zf-CCHH | 14.7 | 117.2 | 19.9 | 3.9 |
| rvp | 16.5 | 133.7 | 55.0 | 11.4 |

#### S5.4 Other experiments

Figure S25 compares methods on simulated datasets without any introduced fragmentation, showing both alignment error and running time; Figure S26 provides a closer look at the runtime comparison between UPP2 and UPP(MAGUS)+adj on these datasets. UPP2, UPP(MAGUS)+adj, MAGUS, and PASTA were the four leading methods across the simulated model conditions. Within this leading group of methods, MAGUS was the most accurate alignment method, closely followed by UPP2, PASTA, and UPP(MAGUS)+adj, in that order. Clustal Omega and T-COFFEE were the least accurate methods. MUSCLE, although more accurate than Clustal Omega and T-COFFEE, was less accurate than the leading group of four methods. UPP2 and MAGUS were the fastest methods, followed by Clustal-Omega and T-COFFEE. UPP(MAGUS)+adj was almost always the slowest, but PASTA, Muscle, and MAFFT linsi were typically also slow. Thus, if selecting from among the most accurate methods, UPP2 and MAGUS are the only ones that provide competitive accuracy and also fast running times.

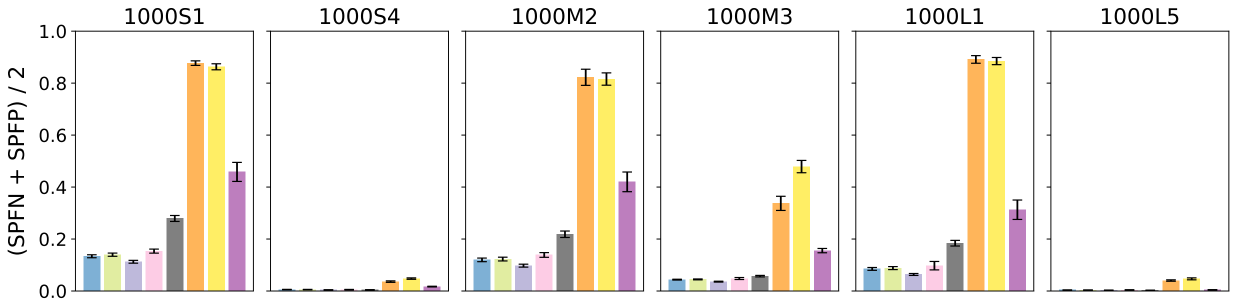

(a) Average alignment error

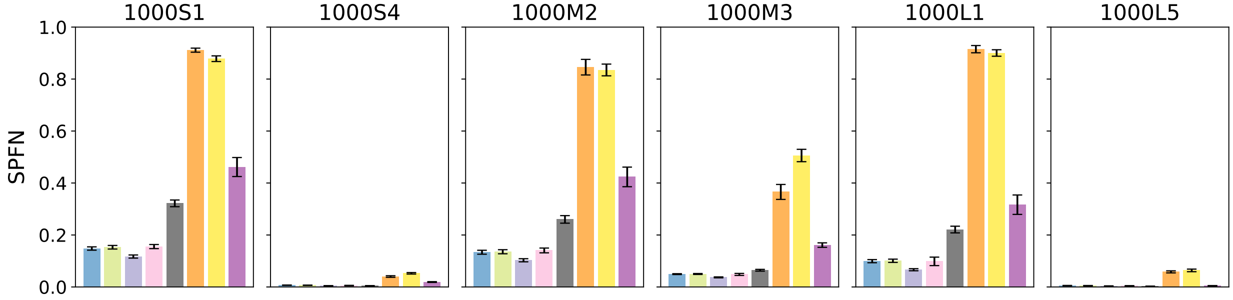

(b) SPFN alignment error

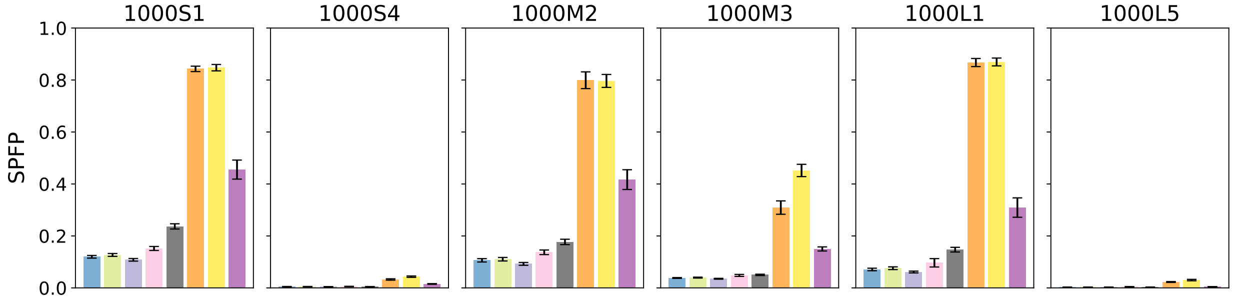

(c) SPFP alignment error

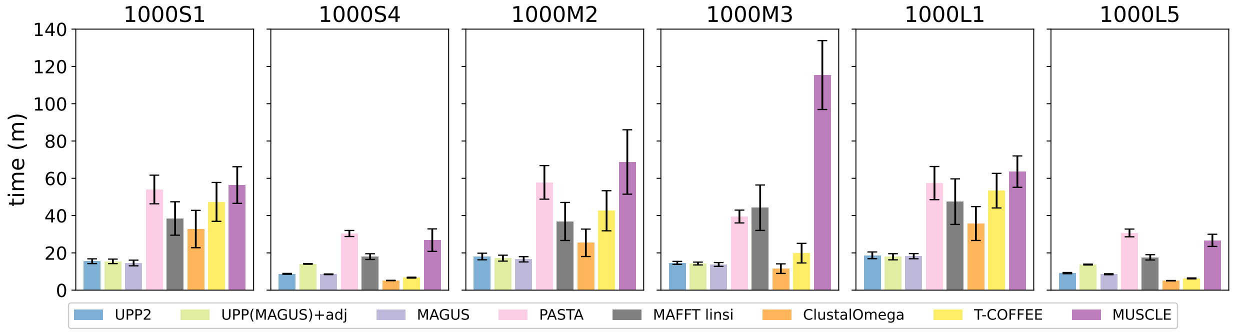

(d) running time

Figure S25: **Experiment 2: UPP2 compared to benchmark alignment methods on simulated datasets without fragmentation.** We show alignment error and runtime of UPP2 (i.e., UPP(MAGUS)+adj+EarlyStop) compared to other alignment methods. All methods except T-COFFEE and MUSCLE were run in their default modes and with 16 threads, when possible. T-COFFEE was run using the regressive mode and MUSCLE was limited to 2 iterations. All datasets have 20 replicates each. The means are shown with error bars indicating standard error for alignment error and standard deviation for running time.

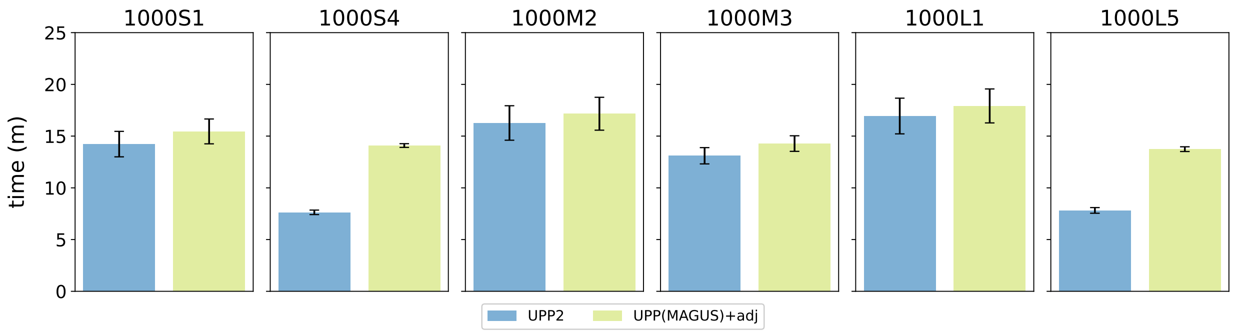

Figure S26: **Experiment 2a: Runtime of UPP2 and UPP(MAGUS)+adj on Simulated Datasets Without Fragmentation** UPP(MAGUS)+adj and UPP2 both use MAGUS backbone alignments, FastTree backbone trees, and adjusted bit-scores; they differ in their search strategies (EarlyStop vs all-against-all). All datasets have 20 replicates each. The means are shown with error bars indicating standard deviation for running time.

### S6 Information about MSA failures

Of the ten Homfam datasets, T-COFFEE failed to run on eight of them due to memory errors. MUSCLE failed to run on the two largest Homfam datasets due to memory errors. We describe the exact errors we encountered below with additional information about the system they ran on.

**MUSCLE failures** MUSCLE had trouble running on the two largest Homfam datasets (zf-CCHH and rvp) with signal 11, also known as segmentation fault.

**T-COFFEE failures** All T-COFFEE runs were done using *T-COFFEE Version_13.45.0.4846264 (2020-10-15 17:52:11 - Revision 5becd5d - Build 620) – regressive mode* in its default mode. They were run on the Illinois Campus Cluster using Singularity. The particular node that the runs were done on had 16 cores available with 128 GB of RAM on a Linux kernel 3.10.0-1160.42.2.el7.x86_64 version #1 SMP Tue Aug 31 20:15:00 UTC 2021. The Singularity image is located at `https://github.com/MinhyukPark/Containers/blob/master/bio.bootstrap`. We ran T-COFFEE after removing the cache directory at $HOME and specifying -cache=no as instructed by the T-COFFEE website, but we were unable to get an output alignment. We also ran T-COFFEE on different nodes on the Campus Cluster, as well as without the Singularity image and installing T-COFFEE, Clustal Omega, and MAFFT locally, but all analyses resulted in the same error.

We also note that on the same system, not even the ROSE datasets could run, resulting in a "free(): double free detected in tcache 2" error. We noticed that by removing sequences from the initial set of sequences, at some point the alignment becomes small enough where T-COFFEE does run without error. These errors and findings were observed with both DNA and Protein sequences.

**Specific T-COFFEE outputs by Homfam dataset** On the PDZ dataset, T-COFFEE fails with "!All Jobs collected" in its output without any indication as to why T-COFFEE was unable to produce any output alignment. The only message indicative of error is "mv: cannot create regular file '0': File exists" in several places in its output.

On the blmb, adh, and Acetyltransf datasets, T-COFFEE fails with "double free or corruption (!prev)" error, which is a typical memory error. It is able to compute the guide tree and weights before segfaulting on computing the MSA step.

On the p450, aat, sdr, zf-CCHH, and rvp datasets, T-COFFEE fails with "double frcee or corruption (!prev)" error during the compute guide tree step.

On the rrm dataset, T-COFFEE fails with "–ERROR: : Impossible to run dynamic.pl" during the compute MSA step.
